# Extensive hybridisation throughout clownfishes evolutionary history

**DOI:** 10.1101/2022.07.08.499304

**Authors:** Sarah Schmid, Baptiste Micheli, Fabio Cortesi, Giulia Donati, Nicolas Salamin

## Abstract

The contribution of hybridisation in the generation of global species diversity has long been controversial among evolutionary biologists. However, it is now increasingly accepted that hybridisation has many impacts on the process of speciation. Notably, it is an important mechanism fostering adaptive radiation since it can generate new phenotypic combinations enabling the occupancy of new niches. Here, we focused on clownfish (Pomacentridae), a clade of 28 coral reef fishes displaying a mutualistic interaction with sea anemones. This behaviour is the key innovation that triggered adaptive radiation of clownfishes, as each species is able to occupy a different combination of host anemone species and habitat. Previous work suggested that hybridisation might be responsible for the extant diversity of clownfish species. To test this hypothesis, we analysed whole-genome datasets for each clownfish species. First, we reconstructed the phylogeny of the clade based on topology weighting methods, which enables the visualisation of the relationships between taxa across the genome. Then, we highlighted possible ancient hybridisation events based on a comparative genomic framework for detecting introgression in genomes. The resulting phylogeny is consistent with previous works based on a few mitochondrial and nuclear genes, and shallow nodes are now well supported in contrast to past studies. Furthermore, we detected multiple past hybridisation events throughout the evolutionary history of clownfishes, corroborating the potential role of hybridisation in the clownfish adaptive radiation. This study adds to the growing number of studies investigating the genomic mechanisms behind species diversification, drawing us closer to understanding how Earth biodiversity is generated.

## INTRODUCTION

At least a tenth of all animal species and more than a quarter of plant species worldwide exhibit evidence of hybridisation – an exchange of genetic material between distinct evolutionary entities (Arnold 1997). Consequently, a wide variety of organisms display mosaic genomes constituted of chromosomal segments from different species or lineages (e.g. Sankararaman et al. 2016, Schumer et al. 2016, Elgvin et al. 2017). Such species thus constitute alluring models to disentangle a broad range of evolutionary processes like the establishment of species barriers or reproductive isolation (Harrison and Larson 2014, Gompert et al. 2017). Indeed, investigating the impact of hybridisation helped highlight crucial regions of the genome involved in reproductive isolation in various species (Schumer et al. 2016, Duranton et al. 2020, Abdelaziz et al. 2021). These efforts have led to a better understanding of the factors maintaining species barriers and how introgression – the incorporation of foreign variants into a local gene pool through backcrossing (Anderson and Hubricht 1938) – can occur between diverging populations (Rafati et al. 2018, Kim et al. 2019). Furthermore, recent development in methods to analyse high-throughput genomic data from non-model organisms open the path to investigate the creative role of hybridisation in the evolution of life on Earth and to focus on the consequence of hybridisation on adaptation and speciation (Seehausen 2004, Abbott et al. 2013, Abbott et al. 2016, Payseur and Rieseberg 2016).

Hybridisation can lead to different processes according to the level of divergence of the two parents. Shortly after population differentiation, when divergence is weak, gene flow between two sister populations will often hinder further diversification due to its homogenization effect on allele frequencies (Weir and Price 2011, Cutter and Gray 2016). However, at an intermediate level of divergence, the genomes are often thought to be only partially permeable to introgression. When exchanges do occur, they might involve genomic regions associated with adaptive traits or neutral regions in linkage with them (Harrison and Larson 2016, Payseur and Rieseberg 2016). The introgression of adaptative alleles into a foreign genomic background might promote diversification through the generation of phenotypic or evolutionary novelties (Arnold and Kunte 2017). Therefore, adaptive introgression has the potential to occur when the degree of divergence between the parental species is within an optimal range. This range is bounded between a minimum divergence required for new and advantageous genotype combinations to evolve and a maximum divergence beyond which the level of genetic incompatibilities is too high, repressing the potential hybridisation benefits (Comeault and Matute 2018). The process of adaptive introgression can lead to various outcomes such as ecological adaptation or expansion of geographic range (Whitney et al. 2010, Jones et al. 2018). Moreover, if adaptive introgression co-occurs with an ecological opportunity, it may provide the high level of heritable genetic diversity required to trigger adaptive radiation (Schluter 2000, Seehausen 2004, Meier et al. 2017, Marques et al. 2019).

Adaptive radiation is characterised by the rapid diversification of an ancestral population into many ecologically diverse species (Schluter 2000). Since the standing genetic variation in a single ancestral population is limited and substantial time is necessary for *de novo* mutations to emerge, previous studies suggested that hybridisation can generate the required genetic variability (Seehausen 2004, Brock and Wagner 2018). Two hypotheses have been put forward to explain the role of hybridisation in adaptive radiation: the *syngameon* and the *hybrid swarm* hypotheses (Seehausen 2004), which differ mainly in the timing of hybridisation and the divergence time between the hybridizing taxa. In the *syngameon* hypothesis, hybridisation occurs after the onset of the adaptive radiation and thus among the members of the same adaptive radiation event, as demonstrated in the *Heliconius* butterflies (Dasmahapatra et al. 2012, Pardo-Diaz et al. 2012) or the Darwin’s finches (Lamichhaney et al. 2015). In contrast, the *hybrid swarm* hypothesis involves hybridisation events occurring before the onset of the adaptive radiation between distantly related species that evolved independently during multiple generations (Seehausen 2004). In this case, the resulting combination of old alleles that was not previously found in neither of the parental species fuel the entire adaptive radiation (Seehausen 2004, Marques et al. 2019). While the combinatorial perspective of speciation offers new insights into our understanding of rapid adaptive radiation events, it has been so far highlighted only in a few non-marine taxa such as in cichlid fishes (Genner and Turner 2012, Meier et al. 2017, Irisarri et al. 2018, Malinsky et al. 2018, Meier et al. 2018, Poelstra et al. 2018), the Hawaiian silversword (Barrier et al. 1999), *Heliconius* butterflies (Wallbank et al. 2016, Enciso-Romero et al. 2017) or Darwin’s finches (Lamichhaney et al. 2015).

Here, we take advantage of the previously described adaptive radiation of the clownfish clades (Litsios et al. 2012) to disentangle the role of ancestral hybridisation events in their rapid diversification. Clownfishes (Pomacentridae subfamily) – also known as anemonefishes – constitute a clade of coral reef fish comprising 28 recognised species and two natural hybrids (Fautin et al. 1997). Their distribution spans from the Indian to the Western Pacific Ocean, with a peak of species richness in the Indo-Malay Archipelago (Elliott and Mariscal 2001). They display an obligate mutualistic interaction with sea anemones – a behaviour considered the key innovation that triggered their adaptive radiation (Litsios et al. 2012). Clownfishes can interact with up to ten sea anemone species, and each clownfish species has a different combination of host anemone and habitat type (Litsios et al. 2012). Direct competition is thus avoided, and various species of clownfish can coexist in a restricted geographical area (Litsios et al. 2012). Previous work has suggested a main hybridisation event at the base of the radiation (Litsios and Salamin 2014), which coincides with a burst of diversification in the clade (Cowman and Bellwood 2011, Frédérich et al. 2013) and may also be linked with an increased rate of morphological evolution (Litsios et al. 2012). Such patterns are consistent with the *syngameon* hypothesis, but the analyses were based on restricted genetic data and only included a fraction of the clownfish species (Litsios and Salamin 2014).

Given this framework, we aimed with this study to further investigate ancestral hybridisation in the clownfish clade and to evaluate the impacts of hybridisation on the genealogical relationships across the clownfishes genome. To achieve this purpose, we first reconstructed the complete phylogeny of the clownfish clade based on whole-genome nuclear SNPs, whole-mitochondria alignment and alignments of all known genes for the 28 clownfish species. The same dataset was then used to detect potential genomic signatures of past introgression. Such analysis might be valuable to potentially highlight parts of the genome exhibiting patterns of adaptive introgression and thus playing a key role in the adaptive radiation process.

## MATERIAL AND METHODS

### Sampling, library preparation, sequencing and read preprocessing

Fin clips (pieces of dorsal fins about 1 cm long) from each of the 28 species of clownfish were collected by different collaborators between 2013 and 2018. We extracted DNA following the Dneasy Blood & Tissue kit standard procedure and we performed twice the final elution in 100 µl of AE buffer (QIAGEN, Hombrechtikon, Switzerland). We quantified the extracted DNA using Qubit® 2.0 Fluorometer (Thermo Fisher Scientific, Waltham, Massachusetts, USA) and evaluated the integrity evaluated by electrophoresis. We followed the TruSeq Nano DNA library prep standard protocol to prepare libraries with a 350 base pair insert size for whole-genome paired-end sequencing (Illumina, San Diego, California, USA). We validated the fragment length distribution of the libraries with a fragment analyzer (Agilent Technologies, Santa Clara, California, USA). Sequencing was performed by the Genomic Technologies Facility of the University of Lausanne and by the Genomic Platform of the University of Geneva. Libraries generated in 2016 were each sequenced on a single lane of Illumina HiSeq 2500 100 paired-end. Libraries prepared in 2017 were pooled by five and each pool was sequenced on a single lane of Illumina HiSeq 2500 100 paired-end. Finally, libraries produced in 2018 were pooled by eight and each pool was sequenced on a single lane of Illumina HiSeq 4000 100 paired-end (see Table S1 for list of samples per year).

After sequencing, we trimmed the raw reads with Trimmomatic v0.36 (Bolger et al. 2014) using the following parameters: ILLUMINACLIP:TruSeq3-PE.fa:2:30:10 || LEADING:3 || TRAILING:3 || SLIDINGWINDOW:4:15 || MINLEN:36. We assessed read quality before and after processing with FastQC v0.11.5 (Andrews 2010).

### Mapping, genotyping and alignment

We mapped the trimmed reads of each individual to the *Amphiprion percula* reference genome (GenBank assembly accession: GCA_003047355.1; assembled in 24 chromosomes; Lehmann et al. 2018) using BWA v0.7.15 (Li and Durbin 2009). Then, we sorted, indexed and filtered the reads according to their mapping quality (>30) using SAMtools v0.1.19. We assigned the reads to a single read-group using Picard Tools v2.9.0 (http://broadinstitute.github.io/picard/), and removed overlapping reads with ATLAS v0.9 (Link et al. 2017). We validated the mapping using various statistics (Bamtools Stats; Picard CollectInsertSizeMetrics; ATLAS BAMDiagnostics). We used ATLAS (Link et al. 2017) to compute genotype likelihood for each sample and to estimate the major and minor alleles. We filtered the resulting VCF file with VCFtools v0.1.15 (Danecek et al. 2011) and only kept sites informative for all samples, with a minimum depth of 2 and a minimum quality of 40. Finally, we used a custom script to reconstruct chromosome-wide alignments based on the generated VCF file (see alignments statistics in Table S3).

### Nuclear phylogenetic reconstruction

We implemented two different approaches to generate chromosome-wide phylogenetic trees. For the first approach, we used the SNP alignments for each chromosome to reconstruct the maximum likelihood trees by applying an ascertainment bias correction as implemented in RAxML v8.2.4 (Stamatakis 2014) (GTR+G model, asc-corr=lewis, 100 bootstraps). For the second approach, we partitioned each chromosome alignment into windows of 5,000 SNPs (total of 13,086 windows). For each window, we reconstructed a maximum likelihood tree with RAxML using the same options as above. We calculated the gene and site concordance factors (gCF and sCF) for each node of each of these phylogenetic trees with IQ-TREE v2.0.5 (Minh et al. 2020). Finally, we inferred a consensus tree for each chromosome based on the previously reconstructed maximum likelihood trees using the multi-species coalescent approach implemented in ASTRAL v5.7.3 (Zhang et al. 2018). We applied the same method to reconstruct a whole-genome nuclear tree based on the 13,086 trees previously inferred with RAxMl..

### Dated phylogeny reconstruction

We reconstructed the calibrated phylogenetic tree of clownfishes based on a subset of the ca. 20,000 previously inferred orthologous genes in 23 Actinopterygii species (10 clownfish and 13 outgroup species; Marcionetti et al. 2019). Due to the computational complexity of inferring a dated phylogenetic tree, we selected the 20 most informative genes. To select those genes, we first reconstructed a phylogenetic tree of the 28 clownfish species for each orthologous gene sequence using RAxML with the same options as described above. Then, we used SortaDate (Smith et al. 2018) to select the 20 most informative genes based on three different criteria: i) genes that have the less topological conflict with a previous inferred species tree; ii) genes that have the lowest variance in the root-to-tip distance; iii) genes that contain a decent amount of molecular evolution (i.e. longest trees indicating more substitutions). We concatenated the alignment of the 20 selected genes and inferred the calibrated phylogenetic tree with BEAST v2.6.2 (Bouckaert et al. 2019). We applied a different GTR+G model of evolution to each partition and ran two parallel analyses, each with 50 million generations, an uncorrelated relaxed clock and assuming a log-normal distribution and a birth-death model for the divergence times. Since no fossil record close to the clownfish group was available, we employed a uniform prior with the bounds set between 10 and 15 Myr to calibrate the root based on a previous estimation of the clownfish divergence times (Litsios et al. 2012). We constrained the topology with the previously defined clownfish clades (Litsios and Salamin 2014), but we estimated the relationships between the clades during the analyses. We set to standard values all other priors. Finally, we summarized the samples of the two runs into a unique dated species tree (posterior probabilities limit of 0.5) with TreeAnnotator v2.6.2 (Bouckaert et al. 2019).

### Mitochondrial phylogenetic reconstruction

We reconstructed the mitochondrial genomes for the 28 different species using MITObim v1.9 (Hahn et al. 2013). We performed an iterative mapping of the trimmed reads to reconstruct the mitochondrial sequence of each clownfish species using the mitochondrial genome of *Amphiprion percula* (GenBank assembly accession: GCA_003047355.2; Lehmann et al. 2018). We also retrieved the whole mitochondrial sequence of one close relative species of the clownfish group, *Pomancentrus moluccensis* (GenBank: MT199209.1). The 29 sequences were aligned with Clustal-Omega v1.2.1 (Sievers and Higgins 2014), producing an alignment of 17,280 nucleotides. To reconstruct the phylogenetic tree, we selected the best model of substitution (i.e. GTR+G+I) with the *phymltest* function (Guindon and Gascuel 2003) from the R package APE v5.3 (Paradis and Schliep 2019). We then performed maximum likelihood reconstruction using RAxML v8.2.4 (Stamatakis 2014) with 100 bootstraps.

### Topology comparison

We compared the tree topologies generated for each of the 24 chromosomes and the whole-genome (total of 25 topologies) by calculating the Kendall-Colijn distance (Kendall and Colijn 2016) between all pairwise combinations using the R package TREEDIST v1.1.1 (Martin R. Smith 2020). We visualised the pairwise distances between the trees using metric multidimensional scaling (MDS; Hillis et al. 2005) as implemented in ADE4 v1.7 (Dray and Dufour 2007). We also investigated the variation in the estimated phylogenetic relationships across the genome. For each of the 5,000 SNPs window (13,086 windows in total), we reconstructed a maximum-likelihood tree with RAxML v8.2.4 (Stamatakis 2014; GTR+G model with a Lewis ascertainment bias correction) and constrained the inference to the main topologies obtained at the chromosome level (seven in total, see Figure 1B). We selected the topology with the best likelihood and calculated the percentage of windows with each of the seven topologies.

**Figure 1.**
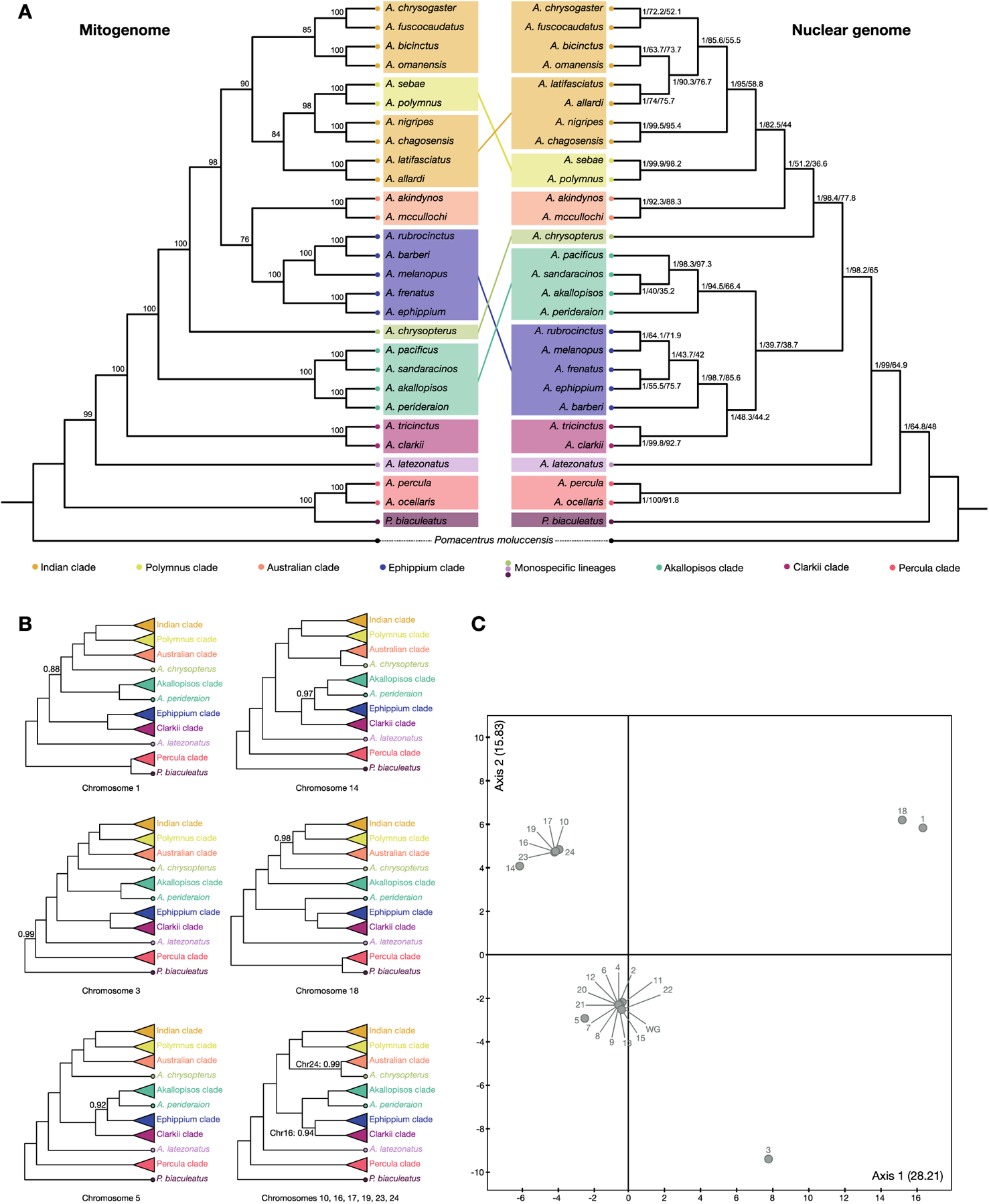
Cladograms of the different genomic partitions of the 28 clownfish species. (A) Comparison of mitochondrial (left) and nuclear (right) phylogenies. Mitochondrial phylogeny was reconstructed based on the whole-mitochondrial sequence using RAxML v8.2.4 and nuclear phylogeny based on whole-genome data for all chromosomes combined (65,378,772 SNPs) using ASTRAL v5.7.3. The dated phylogeny is available in supplementary material (Figure S1). The colors correspond to the different clades. Lines between the two trees indicate different placement of particular clades. Branch lengths are in coalescent unit mutation (i.e. branch length in mutation units divided by the population size parameter) for the nuclear phylogenies and in mean number of substitutions per site for the mitochondrial phylogeny. Node labels of the mitochondrial phylogeny correspond to the bootstrap support based on 100 bootstraps resampling. For the nuclear phylogeny, node labels correspond to the posterior probability / the gene concordance factor (gCF) / the site concordance factor (sCF). A. is for Amphiprion and P. for Premnas. (B) Cladograms showing the relationships for the clownfish clades highlighted above, for each chromosome displaying a different clade topology compared to the species tree. Posterior probability is only indicated for nodes with a value below 1. (C) MDS of pairwise Kendall-Colijn distances between trees reconstructed with Astral for each chromosome (1-24) and for the whole-genome. Eigenvalue of each axis is indicated into bracket.

### Past introgression detection

Considering that the phylogenetic reconstructions showed disparities between both mitochondrial and nuclear trees and among chromosome trees, we assumed that these discrepancies could be due to past introgression events. Therefore, we applied multiple complementary approaches to test whether gene flow could have occurred in the early stages of clownfish diversification. Since we are interested in hybridisation events between the ancestors of the current species, we analyzed the data by grouping species into eleven well-resolved clades, as defined in the reconstructed phylogeny (Figure 1).

#### D statistic, f-4 ratio and f_d_

We tested for historical admixture between all possible combinations of clades using the ABBA-BABA test as well as the related *f*_4_-ratio, as implemented in DSUITE (Malinsky et al. 2019). Both rely on a set of recipient species (referred as P_1_ and P_2_), a donor species (P_3_) and an outgroup (O). The ABBA-BABA test relies on the imbalance between two genealogies to calculate the Patterson’s *D* statistics (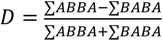, where BABA refers to cases in which P_1_ and P_3_ share the derived allele, and ABBA to cases in which P_2_ and P_3_ share the derived allele), whereas the *f*_4_-ratio is calculated by splitting the potential donor (P_3_) into two subsets according to the following formula 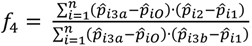. Unlike the *D* statistic, the *f*_4_-ratio estimates the admixture fraction and is comparable between studies. Since both statistics cannot assess introgression between sister lineages, we constricted the possible combinations with the known species topology. We used *Pomacentrus moluccensis* as outgroup taxa (O) and alternatively considered each of the eleven clades as a potential donor (P_3_) or recipient (P_1_ and P_2_). A positive *D* statistic or *f*_4_-ratio suggests introgression between P_3_ and P_2_, whereas a negative value suggests introgression between P_3_ and P_1_. We calculated the statistical significance and standard errors of the resulting *D* statistics and *f*_4_-ratio using a standard block jackknife procedure (Green et al. 2010, Durand et al. 2011) and adjusted the *p*-values for multiple testing using the Benjamini-Hochberg method (Benjamini and Hochberg 1995).

Although the *D* statistic and *f*_4_-ratio can detect introgression genome-wide, they are not appropriate to estimate the proportion of the genome that introgresses in smaller genomic windows (Martin et al. 2015). Therefore, we calculated the *f_d_* statistic to estimate the level of introgression between the P_2_ and P_3_ combinations that had a significant Patterson’s *D* statistic. The *f_d_* statistic (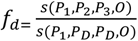 where *s* = ∑(*ABBA* − *BABA*)) is a modified version of the Patterson’s *D* statistic, for which the donor population (P_D_) is defined independently for each site and corresponds to the population (P_2_ or P_3_) with the higher derived allele frequency. We calculated *f_d_* along the genome in non-overlapping windows of 5,000 sites using Martin et al. (2015) scripts.

### Gene ontology and rate of evolution of chromosome 18

Based on the unique topology and introgression signal displayed by chromosome 18, we investigated whether specific gene ontology (GO) terms were enriched in the regions of chromosome 18 showing a strong signal of introgression (from 7,240,000bp to 13,620,000bp and from 28,700,000bp to 34,780,000bp). We used the 17,179 annotated genes inferred by Marcionetti et al. (2019), among which 14,002 were annotated with *biological process* GOs. We retrieved gene annotations for all the genes within the two blocks of interest of chromosome 18 and performed GO enrichment analysis by comparing those annotations to the complete set of protein-coding annotated genes of *A. percula*. We used the R package TOPGO v2.44 (Alexa and Rahnenfuhrer 2020) to perform the gene ontology enrichment analysis based on Fischer’s exact test with the weight01 algorithm. We defined a minimum node size of 10 and considered GO terms with a raw *p*-value below 0.01 as significant, following the recommendations from the TOPGO manual.

Furthermore, we compared the rate of evolution of clownfish genes found in the two blocks of chromosome 18 with a strong introgression signal to the overall rate of evolution by estimating the ratio of non-synonymous over synonymous substitutions (known as dN/dS, Ka/Ks or ω). We expect the dN/dS ratio to be superior to 1 if natural selection promotes gene sequence changes (positive selection) and inferior to 1 if natural selection represses changes (purifying selection). A dN/dS equal to 1 means that the partition is likely evolving neutrally (Kryazhimskiy and Plotkin 2008). We calculated dN/dS values using a perl script (https://github.com/haiwangyang/dNdS). To test whether the mean and median dN/dS values were statistically different in the two blocks of chromosome 18 compared to the rest of the genome, we randomly resampled 10,000 partitions of similar size across the genome using a custom R script. Then, to infer the *p*-value, we compared the mean and median observed dN/dS values to the mean and median values across all resampled partitions.

## RESULTS

### Sequencing, mapping, SNP calling and chromosome alignment statistics

We obtained an average of ca. 169 million raw paired reads across samples, with a number of raw reads per sample ranging from ca. 61 million (*A. mccullochi*) to ca. 534 million (*A. frenatus*; Table S2). After trimming low-quality regions and removing low-quality reads, we ended up with a range of ca. 57 to 482 million paired reads per sample, which corresponds to an estimated raw coverage ranging from 6.4X to 53.6X (Table S2).

We mapped the reads on the *A. percula* reference genome (Lehmann et al. 2018) and obtained between 81% and 97% of reads mapping properly to the reference (i.e. with pairs having the correct orientation and insert-size; Table S3). We proceeded to further filtering by removing low-confidence mapped reads and potentially redundant sequencing data arising from the overlap of paired reads and obtained a final coverage ranging between 4.9X and 40X (Table S3). After mapping, we computed genotype likelihood, estimated the major and minor alleles and filtered the resulting VCF file to obtain a final number of 65,529,334 SNPs.

### Phylogenetic relationships among species and clades

We reconstructed phylogenetic trees based on the whole-mitochondrial genome and whole-genome SNPs to determine the evolutionary history and relationships among the 28 clownfish species. Ancestral nodes for both mitochondrial and nuclear phylogenetic trees are well supported, while more recent splits display lower node support, particularly in the tree inferred with the mitochondrial genome. The two genomic datasets also lead to differences in the topologies obtained. First, the nuclear phylogenetic tree places *Premnas biaculeatus* as the basal lineage to all other clownfishes, while the mitochondrial genome supports a basal group containing the three species *P. biaculeatus*, *A. ocellaris* and *A. percula* (Figure 1). Second, in the nuclear phylogenetic tree, the *clarkii* clade is sister to the *ephippium* clade, and both are clustered with the *akallopisos* clade (Figure 1). However, in the mitochondrial phylogenetic tree, the *clarkii* and the *akallopisos* clades are monophyletic, and the *ephippium* clade is sister to the *Australian* clade (Figure 1). Finally, the *polymnus* clade is interspersed inside the *Indian* clade in the mitochondrial dataset, while the nuclear genome resolves this clade as the sister clade to the whole *Indian* clade (Figure 1A).

Next, to investigate variations in relationships among clownfish clades across chromosomes, we reconstructed consensus phylogenetic trees for each chromosome based on the most likely topology inferred for each genomic window. When considering chromosomes separately, we observe six additional topologies, each showing a specific clade relationship (Figure 1B, 1C). In chromosome 1, *P. biaculeatus* is nested within the *percula* clade, and the *akallopisos* clade is sister to the *Indian*-*polymnus*-*Australian* clades. Chromosome 18 has a similar topology, with the *akallopisos* clade clustering with the *Indian*-*polymnus*-*Australian* clades, except for the species *A. perideraion* which is grouped with the *clarkii* and *ephippium* clades. Chromosome 3 differs from the main topology due to the *akallopisos* clade clustering with the *Indian*-*polymnus*-*Australian* clades instead of the clarkii and ephippium clade. In chromosomes 5 and 14, the *ephippium* clade is sister to the *akallopisos* clade, and chromosome 14 differs from the 5 due to *A. chrysopterus* grouping with the *Australian* clade. Finally, the monospecific lineage *A. chrysopterus* clusters with the *Australian* clade in chromosomes 10, 16, 17, 19, 23 and 24. All other chromosomes display similar clade relationships to the main nuclear consensus topology (Figure S2).

To describe the distribution of the main chromosome topologies across the genome, we inferred the most likely topology in each of the 13,086 windows of 5,000 SNPs defined across the genome. Chromosome 14 topology is the most likely topology in 20.7% of the windows and it is followed by the whole-genome topology (16.7%), chromosome 10 topology (16.2%), chromosome 5 topology (14%) and both chromosome 1 and 3 topologies (9.8% each). Chromosome 18 topology (most likely in 2.9% of the windows) is mainly restricted to two specific parts of chromosome 18 compared to the other six topologies scattered across the genome (Figure 2).

**Figure 2.**
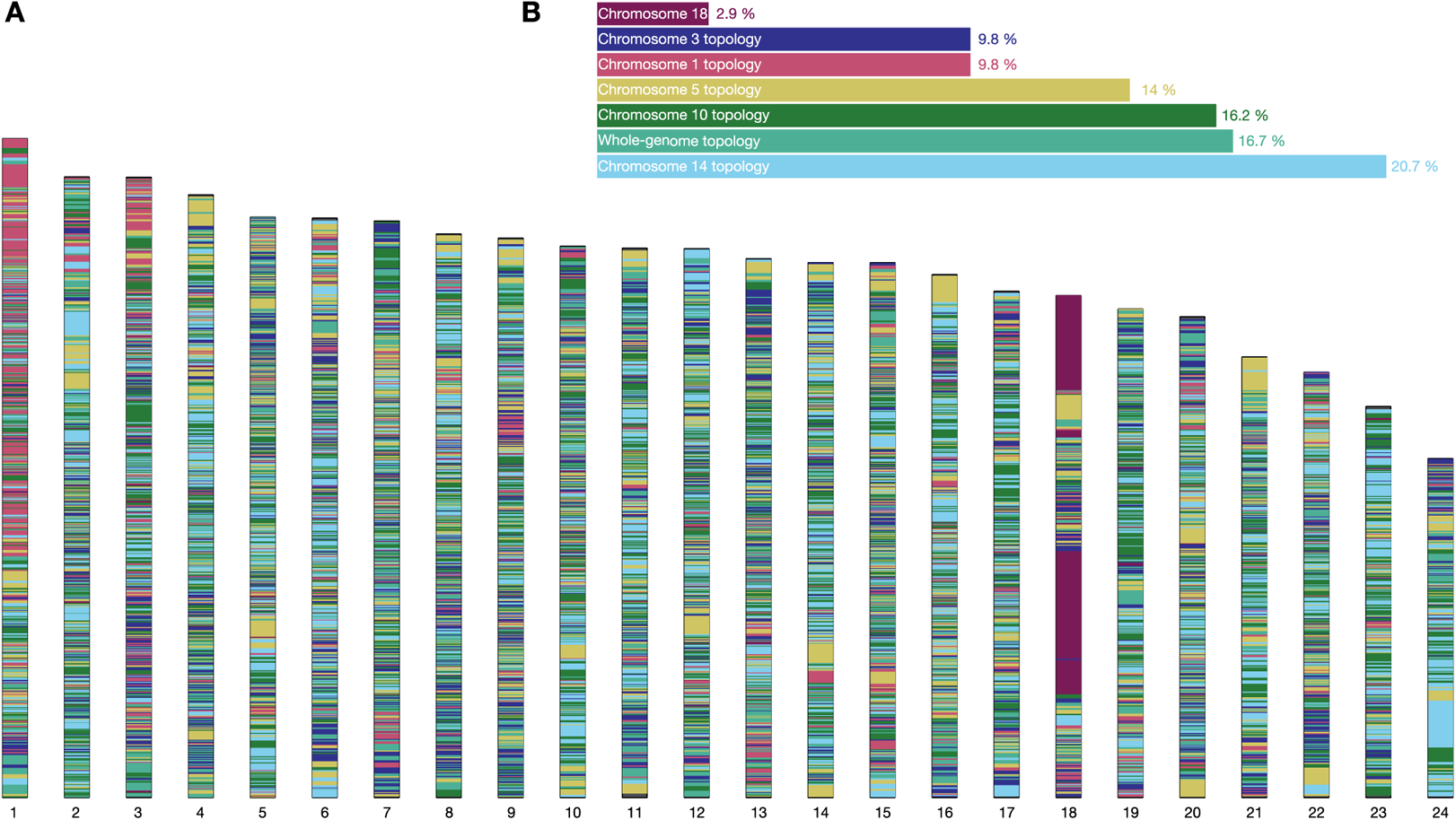
Relationships between the different taxa across the chromosomes. Each of the seven main topologies (see Figure 1) are represented by a colour. (A) For each window of 5,000 SNPs, the topology with the best likelihood is represented. (B) Percentage of windows with a given topology.

Finally, to date the origin of the clownfish radiation and estimate the divergence time among species, we reconstructed a dated phylogenetic tree based on a subset of informative genes. All nodes were highly supported with a Bayesian posterior probability of 1, and the origin of all clownfish clades dates back to 13.21 Myr [95% HPD: 12.73, 13.80] (Figure S1).

### Past introgression among clownfish clades

To investigate the timing and strength of introgression during clownfish adaptive radiation, we compared several independent approaches. The different methods all highlighted past introgression among clownfish species and clades. First, the discordances among both well supported mitochondrial and nuclear phylogenies suggest several events of ancestral hybridisation among clownfish species. Then, we used *D* statistics and *ff*_4_-ratio to quantify the potential widespread introgression among the different clownfish clades (Figure 3, Table S1). We found significant *D* statistics between the three most ancestral lineages of clownfish (*i.e. Premnas biaculeatus*, *A. percula* clade and *Amphiprion latezonatus*) and each one of the most recent clades (range of *D* statistics: 0.003-0.153; Figure 3A). The *ff*_4_-ratio varies between 0.0008 and 0.024 across chromosomes, except for chromosome 18 which displays *ff*_4_-ratio up to four times higher for introgression events involving *A. latezonatus* and the members of the *akallopisos-A. perideraion-ephippium-clarkii* clade. Introgression also occurs among members of most recent clownfish clades at a similar level for all chromosomes (Figure 3B, Table S1), except for chromosome 18, which displays a strong introgression signal between the members of the *Indian*-*polymnus*-*Australian-A. chrysopterus* clade and the *akallopisos* clade as well as between the *ephippium-clarkii* clade and *A. perideraion*, with *ff*_4_-ratio reaching values of 0.5. The high level of introgression observed in chromosome 18 is constricted to two main portions of the chromosome (from ca. 7.24 Mb to 13.62 Mb and from 28.7 Mb to 34.78 Mb; Figure 3C).

**Figure 3.**
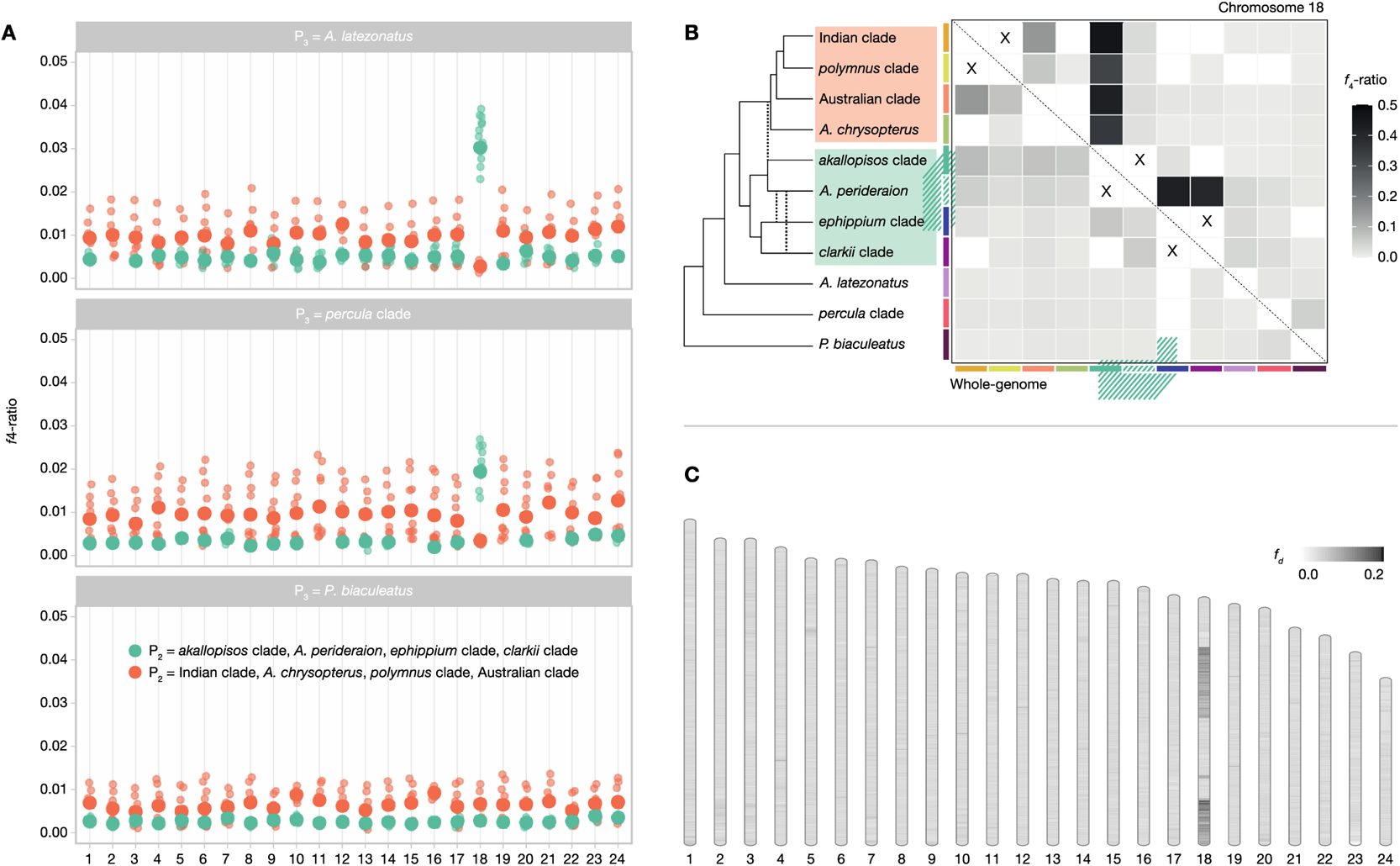
Pattern of hybridisation among clades across the clownfish genome. (A) Significant *ff*_4_-ratio after Benjamini-Hochberg correction for each chromosome considering only the three more ancestral clades/species as P_3_. In turquoise is highlighted introgression between the members of the following clades/species considered as P_2_: Akallopisos clade, *A. perideraion*, Ephippium clade or Clarkii clade and P_1_ consists either in the Indian clade, *A. chrysopterus*, Polymnus clade or Australian clade. In orange is highlighted introgression between P_3_ and one of the following clades/species considered as P_2_: *A. chrysopterus*, Polymnus clade or Australian clade. Big dots correspond to the mean *ff*_4_-ratio and small dots are the values for each combination. (B) Introgression between all the different clades/species. The lower triangle represents the *ff*_4_-ratio values calculated over the whole-genome, while the upper triangle represents the *ff*_4_-ratio values calculated only over the chromosome 18. Each grey square stand for a specific combination of P_2_ (left of the square) and P_3_ (bottom of the square). Introgression between sister taxa cannot be assessed and are marked by a x. Dashed line on the cladogram represent the strong introgression events highlighted in chromosome 18. The clades highlighted in orange and blue in the cladogram correspond to the clades highlighted in Figure 1A. (C) *f_d_* values across the 24 chromosomes calculated in 5,000 SNPs windows and average among all combinations of P_2_ and P_3_. (see Supp. Mat. for the precise *ff*_4_-ratio, *D* statistic, *f_d_* and *p*-values for each combination and chromosome).

### Chromosome 18

To investigate the potential reasons for the elevated introgression and unique topology of the two blocks of chromosome 18, we proceeded to gene ontology (GO) enrichment and calculated the dN/dS ratio of the genes within those regions. We identified 361 annotated genes in the two blocks of chromosome 18 and highlighted 20 significant enriched terms (*p*-value < 0.01; Tables S7 and S8). Among them, we found GO terms associated with DNA damage response mediated by p53 (GO:0043518, GO:1902254, GO:0043517), various processes linked with the nervous system (GO:0035633, GO:0090249, GO:0021530, GO:0014044, GO:0021508), phospholipids metabolic process (GO:0006644), processes related to embryonic development (GO:0090191, GO:0007525, GO:0003337, GO:0060712) and responses to external stressors (GO:0071480, GO:0002438, GO:0038096; see Table S8 for the complete list). Finally, the mean (0.20) and median (0.16) dN/dS of the two blocks of chromosome 18 were not significantly different from the ones estimated by resampling across all the whole-genome (Table S9, Figure S3).

## DISCUSSION

The present work is part of the growing number of genomic studies on hybridisation between related species illustrating that this phenomenon is widespread across the tree of life. The description of the phylogenetic relationship of the 28 clownfish species associated with investigating past introgression events highlighted widespread hybridisation among clownfish species, with some parts of the genome showing a strong introgression signal. This study contributes to a better understanding of all the genetic and ecological mechanisms at play during the process of adaptive radiation.

### Phylogeny and cytonuclear discordances

We reconstructed a comprehensive phylogenetic tree for the 28 known clownfish species – based on both complete nuclear and mitochondrial genomes – which updates considerably previous inferences relying on few genes and provides a better understanding of the evolutionary history of all clownfishes. The nuclear and mitochondrial clades identified in this study are consistent with those previously described (Litsios and Salamin 2014), except for the *percula* clade at the nuclear level, which encompasses only *A. ocellaris* and *A. percula*, thus making *P. biaculeatus* a monophyletic clade. Cytonuclear discordances at the clade level were also previously revealed. In the nuclear phylogeny, the *clarkii* clade is sister to the *akallopisos* and *ephippium* clades, a relationship disappearing in the mitochondrial phylogeny. Both nodes in the mitochondrial and nuclear phylogenies were strongly supported, suggesting that the observed discordances can be a consequence of either incomplete lineage sorting or hybridisation. Incomplete lineage sorting – the process leading to the persistence of ancestral polymorphisms across successive speciation events – was documented in various groups of organisms across the Tree of Life (e.g. Wang et al. 2018, Lopes et al. 2021, Meleshko et al. 2021, Dong et al. 2022, Feng et al. 2022) and is known to be difficult to disentangle from hybridisation (Joly et al. 2009, Yu et al. 2013). In the context of adaptive radiations, cytonuclear discordance between basal branches are commonplace (reviewed in Seehausen 2004) and often coincide with hybridisation predating the radiation. Indeed, hybridisation can increase the amount of standing genetic variation, subsequently creating adaptive novelty, which is a key mechanism in adaptive radiation (Grant and Grant 1997, Seehausen 2004). In clownfish, hybridisation is frequent in captivity and two recognised hybrid species are known in the wild: *A. leucokranos* and *A. thiellei* (Fautin et al. 1997). Furthermore, additional genomic evidence (see Discussion below) points toward hybridisation rather than incomplete lineage sorting as the source of cytonuclear discordance.

### A variety of chromosome and windows topologies

Although the phylogeny mentioned above displays the maximal likelihood when using whole-genome SNP data, it does not represent the complex relationship between the different species. Thus, chromosome-wide and windows-based phylogenies were also reconstructed, offering a fine-scale perspective of the relationship between species across the genome. The windows-based approach highlighted a wide array of topologies. Surprisingly, when constricting the phylogenetic reconstruction to the seven main topologies and selecting the most likely topology for each window, the whole-genome topology was in the second position after the chromosome 14 topology in terms of percentage of windows. A possible explanation for this contradictory result is that the whole-genome topology might often be ranked as one of the most likely topologies in multiple windows (but not the most likely). In contrast, the chromosome 14 topology, outside of its most likely windows, is not highly ranked in other windows. Such conflicts between the different partitions and chromosomes were long considered a nuisance. However, recent works acknowledged that those patterns contain helpful information about the diversification process (Marin et al. 2020). Indeed, gene duplication, incomplete lineage sorting, introgression and hybridisation could produce mosaic genomes made of segments with different evolutionary histories (Salzburger 2018). Interestingly, numerous genomes from species arising from adaptive radiation display mosaic patterns (Mallet et al. 2007, Brawand et al. 2014, Fontaine et al. 2015), such as the cichlid fishes genome, which consists of more than two thousand distinct topologies (Salzburger 2018).

### Widespread introgression and past hybridisation

The described cytonuclear discordances and conflicting topologies among chromosomes and genome partitions suggest recurrent hybridisation events during the evolutionary history of the clownfish group. We revealed further evidence of hybridisation using the *D* statistics and *f*_4_-ratio, which highlight a widespread signal of hybridisation among all clownfish clades. Similar to clownfishes, low introgression among clades and across chromosomes (except for chromosome 18, see discussion below) was also found in various species displaying signatures of past introgression (Barth et al. 2020, Svardal et al. 2020). Different factors such as phylogeny, geographical distance and the species’ biology can influence the frequency of hybridisation (Hamlin et al. 2020). Indeed, we expect more genetic exchange between recently diverged taxa (Coyne and Orr 1997) as well as between species with external fertilization and without sexual chromosomes (Runemark et al. 2019), such as clownfishes.

More specifically, we highlighted introgression between the three most ancestral clades and the other younger clades. This result suggests that events of ancestral hybridisation might have taken place at the base of the clownfish radiation. Hybridisation occurring at the onset of adaptive radiation was already described in various species, such as the African cichlids (Meier et al. 2017). Furthermore, it was recently suggested that hybridisation could reshuffle old genetic variations into new combinations eventually leading to speciation or adaptive radiation (Marques et al. 2019). Contrary to new mutations, old variations have already been through a selection filter and fit their genomic and ecological context (Abbott et al. 2013). Several examples of species arising from new combinations of old alleles exist (see Marques et al. 2019 for a review), and many of them are well-known cases of adaptive radiations. Clownfish are thus well-suited candidates for this “combinatorial view of speciation”, but further evidence is still needed to establish a causal link between past hybridisation events and the diversification of clownfishes. The identification of such causal link could be achieved by comparing of the age of the ancient adaptive alleles and the species splitting time, combined to the estimation of linkage disequilibrium patterns of such alleles. However, further knowledge of the genes involved in the clownfish radiation is required to perform such analyses.

Additionally, we cannot rule out the possibility of one or several introgression events with an unsampled or extinct ancestral species. Such events known as ghost introgression have the potential to enhance adaptation and speciation (Ottenburghs 2020). Considering archaic introgressed tract opens new horizons to disentangle the impact of those ancestral variations on the recipient lineage, thus improving our understanding of the role of introgression in the evolutionary trajectory of species (Jacobs and Therkildsen 2019). However, up to now, only a few studies have investigated in detail archaic tracts in non-hominid genomes and focused, among others, on the sea bass (Duranton et al. 2019), the killer whale (Foote et al. 2019) and the bonobo (Kuhlwilm et al. 2019).

Additionally, the extent of introgression between clownfish species across the genome does not provide information on whether the process is mainly neutral or promoted by a selective advantage. It was previously reported that clownfishes’ rapid radiation coincides with their mutualism with sea anemone (Litsios et al. 2012). Nevertheless, knowledge about the genetic basis of the mutualism between the clownfish and the sea anemone is still at an early stage. Seventeen positively selected genes at the basis of the clownfish radiation were previously highlighted, some coding for functions associated with *N*-acetylated sugars (Marcionetti et al. 2019). These molecules are known to play a role in sea anemone discharge of toxins (Marcionetti et al. 2019). However, the causal link between those genes and the ability to interact with the sea anemone should be validated with other experimental approaches since an improved understanding of the genetic basis underlying this mutualism interaction is essential for identifying adaptive introgressed segments in the clownfish genome.

### The special case of chromosome 18

We highlighted distinct phylogenetic and introgression patterns in chromosome 18 compared to the rest of the genome. Indeed, two large blocks of the chromosome (from ca. 7.24 Mb to 13.62 Mb and from 28.7 Mb to 34.78 Mb; Figure 3C) display strong introgression signals among some clownfish clades, which presumably led to the unique topology exhibited by those two regions. The maintenance of the two blocks in chromosome 18 could be explained by structural variations such as inversions, which might prevent recombination within these regions (Kirkpatrick 2010, Stevison et al. 2011). Inversions are known to suppress recombination locally and are responsible for the persistence of supergenes in various species (Joron et al. 2011, Li et al. 2016, Tuttle et al. 2016). In comparison, recombination is probably still ongoing in the rest of the genome and generates – in combination with introgression – the mosaic of topologies we observed. Similar to this work, unusually long blocks containing minor topologies were also described in *Heliconius* butterflies and were suggested to be inversions captured by introgression (Edelman et al. 2019). Although the evolution of sex chromosomes could produce similar chromosome patterns (Natri et al. 2019), it is an unlikely scenario in the clownfishes. Indeed, the 28 species are sequential hermaphrodites and thus do not exhibit sex chromosomes (Arai and Inoue 1976, Fricke 1979) but instead bear genes related to sex change spread throughout the genome (Casas et al. 2018)

Although further investigation is required to confirm the presence of structural variations within chromosome 18, the high level of introgression in these two blocks compared to the rest of the genome suggests that hybridisation played a role in the maintenance of these regions. Furthermore, the two portions of chromosome 18 encompass genes linked with processes such as DNA damage and external stressors responses, nervous system and embryonic development, as well as phospholipids metabolism. These functions could be linked to advantageous traits involved in local adaptation and pre- or postzygotic isolation, as suggested within inverted regions of various species(Kirkpatrick and Barton 2006, Lowry and Willis 2010, Ayala et al. 2012).

## Conclusion

This work is part of the growing body of evidence showing that hybridisation is an important evolutionary force not only in plants but also in animals (Mallet 2007, Abbott et al. 2013). Here, we used the *D* statistic and the *f_4_*-ratio as introgression estimators. Although they have the advantage of being simple to use, they are nevertheless limited in terms of information about the timing and the direction of introgression (Martin et al. 2015). In addition, our study also questions the use of bifurcating diagrams since it seems that a single tree-like phylogeny does not capture the clownfishes’ complete and complex evolutionary history.

Despite these limitations, the cytonuclear discordance, the conflicting topology between the different partitions of the genome, the *f*_4_-ratio and the *D* statistic as well as the known occurrences of current clownfish hybrid species all point toward multiple past introgression events, which might have fuelled the adaptive radiation of clownfish. A direct causal link remains to be established by finding a functional role of the past introgression of this intriguing case of adaptive radiation.

## AUTHOR CONTRIBUTIONS

NS and SS planned the study. FC and GD collected part of the samples. BM and SS performed lab work and analysed the data. SS wrote the manuscript. All authors contributed to the final version of the manuscript.

## ACKNOWLEDGEMENTS

We gratefully thank Glenn Litsios for the collection of most samples used in this study and to Serges Planes for contributing with one sample. We also thank Diego Hartasánchez for the insightful comments on the manuscript and Anna Marcionetti and Joris Bertrand for the fruitful discussions.

## SUPPORTING INFORMATIONS

**Figure S1**. Dated phylogeny of the 28 clownfish species based on the 20 most informative genes using BEAST v2.6.2. Error bars represent the 95% HPD (height posterior density) intervals for the divergence date estimates. Error bars on nodes show the dating confidence interval. Time scale is given in million years before present.. The Bayesian posterior probability is indicated on the right of each node.

**Figure S2**. Phylogeny for each chromosome of the 28 clownfish species based on SNPs using RAxML v8.2.4 and ASTRAL v5.7.3. Node labels correspond to the posterior probability / the gene concordance factor (gCF) / the site concordance factor (sCF). A. is for *Amphiprion* and P. for *Premnas*.

**Figure S3**. Mean and median dN/dS. (A) Histogram of the mean dN/dS across resampled whole-genome windows. The pink dashed lines correspond to the 5% and 95% quantiles of the resampled distribution. The pink line is the mean dN/dS across the 10,000 resampled windows and the blue line is the mean observed dN/dS value in the two partitions of chromosome 18 with high level of introgression and unique topology. (B) Histogram of the median dN/dS across resampled whole-genome windows. The pink dashed lines correspond to the 5% and 95% quantiles of the resampled distribution. The pink line is the median dN/dS across the 10,000 resampled windows and the blue line is the median observed dN/dS value in the two partitions of chromosome 18 with high level of introgression and unique topology.

**Table S1**. Sampling information for each individual used in this project

**Table S2**. Sequencing and read processing statistics. Number of paired reads (sum of reverse and forward reads) after sequencing and after quality filtering. We estimated the coverage by multiplying the number of raw reads with the read length, and divided it by the approximate reference genome size (900 Mb)

**Table S3**. Mapping statistics. Number and percentage of total reads and proper-paired reads mapping to *Amphiprion percula* reference genome (Lehman *et al*. 2018). Properly paired means that paired-reads (forward and reverse) orientation is correct and that the gap between them corresponds to the expected insert size (~350 pb). The number of final mapped reads correspond to the number of reads after filtering and that were actually use for all subsequent analysis. We estimated the coverage by multiplying the number of final mapped reads with the read length, and divided it by the approximate reference genome size (900 Mb)

**Table S4**. Alignment statistics. Statistics for chromosome-wide alignments reconstructed using custom scripts. The number of genes is based on the genes inferred by Marcionetti et al. 2019.

**Table S5**. Introgression statistics (*D*-statistics and *f4*-ratio) among clownfish clades. *D*-statistics and *f4*-ratio for all combinations compatible with the inferred phylogeny for all chromosomes individually (1-24) and for the whole-genome (WG), based on SNP data. The outgroup is always *Pomacentrus moluccensis*. *p*-values are adjusted for multiple comparisons. ABBA: number of sites at which P2 and P3 species/clade share the derived allele. BABA: number of sites at which species/clade P1 and P3 share the derived allele.

**Table S6**. *f_d_* statistics across the genome. *F_d_* statistics calculated across the genome in non-overlapping 5,000 SNPs windows using Martin et al. (2015) scripts. Chromosome, start [bp], end [bp] and *f_d_* value of each window for each combination of clades are indicated. Based on the eleven clades previously defined. aka: *akallopisos* clade (*A. akallopisos, A. pacificus, A. sandaracinos*); aus: Australian clade (*A. akindynos, A. mccullochi*); chp: *A. chrysopterus* (monospecific); cla: *clarkii* clade (*A. clarkii, A. tricinctus*); eph: *ephippium* clade (*A. barberi, A. epphipum, A. frenatus, A. melanopus, A. rubrocinctus*); ind: Indian clade (*A. bicinctus, A. chrysogaster, A. fuscocaudatus, A. omanensis*); ltz: *A. latezonatus* (monospecific); pol: polymnus clade (*A. polymnus, A. sebae*); pbi: *P. biaculeatus* (monospecific); prc: percula clade (*A. ocellaris, A. percula*); prd: *A. perideraion* (monospecific). For example: “aka_aus_ltz” means that P1 is the akallopisos clade, P2 is the Australian clade and P3 is *A. latezonatus*.

**Table S7**. Annotation of the genes in the chromosome 18 partitions with high level of introgression and displaying an alternative topology. For each ENSEMBL gene ID, we provide the chromosome where it is located, the start and end positions of the gene, the external gene name, and the gene ontology (GO) annotation.

**Table S8**. GO enrichment of chromosome 18. Significant gene ontology (GO) terms based on the *weight01* algorithm using a classic-Fischer test to calculate the *p*-value. All significant GO terms (*p* <0.01) found in the two chromosome 18 partitions with high level of introgression and displaying an alternative topology are listed. Description of each term, number of annotated genes across the whole-genome, number of significant genes, number of expected genes and classic-Fischer *p*-value are indicated for each GO term.

**Table S9**. dn/ds value of chromosome 18 windows with high introgression level and unique topology. List of genes inferred by Marcionetti et al. (2019) wound in the windows of chromosome 18 with high introgression level and displaying a unique topology. Start, end, dn/ds value and gene length are indicated for each gene.

## REFERENCES

Abbott, R.J., Albach, D., Ansell, S., Arntzen, J.W., Baird, S.J.E., Bierne, N., Boughman, J., Brelsford, A., Buerkle, C.A. & Buggs, R. (2013) Hybridization and speciation. Journal of Evolutionary Biology 26, 229– 246.

Abbott, R.J., Barton, N.H. & Good, J.M. (2016) Genomics of hybridization and its evolutionary consequences. Molecular Ecology 25, 2325–2332. https://doi.org/10.1111/mec.13685

Abdelaziz, M., Muñoz-Pajares, A.J., Berbel, M., García-Muñoz, A., Gómez, J.M. & Perfectti, F. (2021) Asymmetric Reproductive Barriers and Gene Flow Promote the Rise of a Stable Hybrid Zone in the Mediterranean High Mountain. Frontiers in Plant Science 12.

Alexa, A. & Rahnenfuhrer, J. (2020) topGO: Enrichment analysis for Gene Ontology. R package version 2.43.0.

Anderson, E. & Hubricht, L. (1938) Hybridization in Tradescantia. Iii. the Evidence for Introgressive Hybridization. American Journal of Botany 25, 396–402. https://doi.org/10.1002/j.1537-2197.1938.tb09237.x

Andrews, S. (2010) FastQC: a quality control tool for high throughput sequence data. Available online at: http://www.bioinformatics.babraham.ac.uk/projects/fastqc/. http://www.bioinformatics.babraham.ac.uk/projects/fastqc/

Arai, R. & Inoue, M. (1976) Chromosomes of seven species of Pomacentridae and two species of Acanthuridae from Japan. Bulletin of the National Museum of Nature and Science, Series A, Zoology 2, 73–78.

Arnold, M.L. (1997) Natural Hybridization and Evolution. Oxford University Press, Oxford, New York.

Arnold, M.L. & Kunte, K. (2017) Adaptive Genetic Exchange: A Tangled History of Admixture and Evolutionary Innovation. Trends in Ecology & Evolution 32, 601–611. https://doi.org/10.1016/j.tree.2017.05.007

Ayala, D., Guerrero, R.F. & Kirkpatrick, M. (2012) Reproductive Isolation and Local Adaptation Quantified for a Chromosome Inversion in a Malaria Mosquito. Evolution 67, 946–958.

Barrier, M., Baldwin, B.G., Robichaux, R.H. & Purugganan, M.D. (1999) Interspecific hybrid ancestry of a plant adaptive radiation: allopolyploidy of the Hawaiian silversword alliance (Asteraceae) inferred from floral homeotic gene duplications. Molecular Biology and Evolution 16, 1105–1113. https://doi.org/10.1093/oxfordjournals.molbev.a026200

Barth, J.M.I., Gubili, C., Matschiner, M., et al. (2020) Stable species boundaries despite ten million years of hybridization in tropical eels. Nature Communications 11, 1–13. https://doi.org/10.1038/s41467-020-15099-x

Benjamini, Y. & Hochberg, Y. (1995) Controlling the False Discovery Rate: A Practical and Powerful Approach to Multiple Testing. Journal of the Royal Statistical Society. Series B (Methodological) 57, 289–300.

Bolger, A.M., Lohse, M. & Usadel, B. (2014) Trimmomatic: a flexible trimmer for Illumina sequence data. Bioinformatics 30, 2114–2120. https://doi.org/10.1093/bioinformatics/btu170

Bouckaert, R., Vaughan, T.G., Barido-Sottani, J., et al. (2019) BEAST 2.5: An advanced software platform for Bayesian evolutionary analysis. PLOS Computational Biology 15, e1006650. https://doi.org/10.1371/journal.pcbi.1006650

Brawand, D., Wagner, C.E., Li, Y.I., et al. (2014) The genomic substrate for adaptive radiation in African cichlid fish. Nature 513, 375. https://doi.org/10.1038/nature13726

Brock, C.D. & Wagner, C.E. (2018) The smelly path to sympatric speciation? Molecular Ecology 27, 4153–4156. https://doi.org/10.1111/mec.14845

Casas, L., Saenz-Agudelo, P. & Irigoien, X. (2018) High-Throughput Sequencing and Linkage Mapping of a Clownfish Genome Provide Insights on the Distribution of Molecular Players Involved in Sex Change. Scientific Reports 8, 4073. https://doi.org/10.1038/s41598-018-22282-0

Cowman, P. & Bellwood, D. (2011) Coral reefs as drivers of cladogenesis: expanding coral reefs, cryptic extinction events, and the development of biodiversity hotspots. Journal of evolutionary biology. https://doi.org/10.1111/j.1420-9101.2011.02391.x

Coyne, J.A. & Orr, H.A. (1997) “Patterns of Speciation in Drosophila” Revisited. Evolution 51, 295–303. https://doi.org/10.1111/j.1558-5646.1997.tb02412.x

Cutter, A.D. & Gray, J.C. (2016) Ephemeral ecological speciation and the latitudinal biodiversity gradient. Evolution 70, 2171–2185. https://doi.org/10.1111/evo.13030

Danecek, P., Auton, A., Abecasis, G., et al. (2011) The variant call format and VCFtools. Bioinformatics 27, 2156–2158. https://doi.org/10.1093/bioinformatics/btr330

Dasmahapatra, K.K., Walters, J.R., Briscoe, A.D., et al. (2012) Butterfly genome reveals promiscuous exchange of mimicry adaptations among species. Nature 487, 94–98. https://doi.org/10.1038/nature11041

Dong, W., Liu, Y., Li, E., Xu, C., Sun, J., Li, W., Zhou, S., Zhang, Z. & Suo, Z. (2022) Phylogenomics and biogeography of Catalpa (Bignoniaceae) reveal incomplete lineage sorting and three dispersal events. Molecular Phylogenetics and Evolution 166, 107330. https://doi.org/10.1016/j.ympev.2021.107330

Dray, S. & Dufour, A.-B. (2007) The ade4 Package: Implementing the Duality Diagram for Ecologists. Journal of Statistical Software 22. https://doi.org/10.18637/jss.v022.i04

Durand, E.Y., Patterson, N., Reich, D. & Slatkin, M. (2011) Testing for Ancient Admixture between Closely Related Populations. Molecular Biology and Evolution 28, 2239–2252. https://doi.org/10.1093/molbev/msr048

Duranton, M., Allal, F., Valière, S., Bouchez, O., Bonhomme, F. & Gagnaire, P. (2020) The contribution of ancient admixture to reproductive isolation between European sea bass lineages. Evolution Letters. https://doi.org/10.1002/evl3.169

Duranton, M., Allal, F., Valière, S., Bouchez, O., Bonhomme, F. & Gagnaire, P.-A. (2019) The contribution of ancient admixture to reproductive isolation between European sea bass lineages. Evolutionary Biology. https://doi.org/10.1101/641829

Elgvin, T.O., Trier, C.N., Tørresen, O.K., Hagen, I.J., Lien, S., Nederbragt, A.J., Ravinet, M., Jensen, H. & Sætre, G.-P. (2017) The genomic mosaicism of hybrid speciation. Science Advances 3, e1602996. https://doi.org/10.1126/sciadv.1602996

Enciso-Romero, J., Pardo-Díaz, C., Martin, S.H., Arias, C.F., Linares, M., McMillan, W.O., Jiggins, C.D. & Salazar, C. (2017) Evolution of novel mimicry rings facilitated by adaptive introgression in tropical butterflies. Molecular Ecology 26, 5160–5172. https://doi.org/10.1111/mec.14277

Fautin, D.G., Allen, G.R. & Fautin, D. (1997) Field Guide to Anemone Fishes and Their Host Sea Anemones. 2nd Revised Edition.,. Western Australian Museum.

Feng, S., Bai, M., Rivas-González, I., et al. (2022) Incomplete lineage sorting and phenotypic evolution in marsupials. Cell 185, 1646–1660.e18. https://doi.org/10.1016/j.cell.2022.03.034

Fontaine, M.C., Pease, J.B., Steele, A., et al. (2015) Extensive introgression in a malaria vector species complex revealed by phylogenomics. Science 347, 1258524. https://doi.org/10.1126/science.1258524

Foote, A.D., Martin, M.D., Louis, M., et al. (2019) Killer whale genomes reveal a complex history of recurrent admixture and vicariance. Molecular Ecology 0, 1–18. https://doi.org/10.1111/mec.15099

Frédérich, B., Sorenson, L., Santini, F., Slater, G.J. & Alfaro, M.E. (2013) Iterative Ecological Radiation and Convergence during the Evolutionary History of Damselfishes (Pomacentridae). The American Naturalist 181, 94–113. https://doi.org/10.1086/668599

Fricke, H.W. (1979) Mating System, Resource Defence and Sex Change in the Anemonefish Amphiprion akallopisos. Zeitschrift für Tierpsychologie 50, 313–326. https://doi.org/10.1111/j.1439-0310.1979.tb01034.x

Genner, M.J. & Turner, G.F. (2012) Ancient Hybridization and Phenotypic Novelty within Lake Malawi’s Cichlid Fish Radiation. Molecular Biology and Evolution 29, 195–206. https://doi.org/10.1093/molbev/msr183

Gompert, Z., Mandeville, E.G. & Buerkle, C.A. (2017) Analysis of Population Genomic Data from Hybrid Zones. Annual Review of Ecology, Evolution, and Systematics 48, 207–229. https://doi.org/10.1146/annurev-ecolsys-110316-022652

Grant, P.R. & Grant, B.R. (1997) Hybridization, Sexual Imprinting, and Mate Choice. The American Naturalist 149, 1–28.

Green, R.E., Krause, J., Briggs, A.W., et al. (2010) A Draft Sequence of the Neandertal Genome. Science 328, 710–722. https://doi.org/10.1126/science.1188021

Guindon, S. & Gascuel, O. (2003) A simple, fast, and accurate algorithm to estimate large phylogenies by maximum likelihood. Systematic Biology 52, 696–704. https://doi.org/10.1080/10635150390235520

Hahn, C., Bachmann, L. & Chevreux, B. (2013) Reconstructing mitochondrial genomes directly from genomic next-generation sequencing reads—a baiting and iterative mapping approach. Nucleic acids research 41, e129–e129.

Hamlin, J.A.P., Hibbins, M.S. & Moyle, L.C. (2020) Assessing biological factors affecting postspeciation introgression. Evolution Letters n/a. https://doi.org/10.1002/evl3.159

Harrison, R.G. & Larson, E.L. (2016) Heterogeneous genome divergence, differential introgression, and the origin and structure of hybrid zones. Molecular ecology.

Harrison, R.G. & Larson, E.L. (2014) Hybridization, Introgression, and the Nature of Species Boundaries. Journal of Heredity 105, 795–809.

Hillis, D.M., Heath, T.A. & John, K.St. (2005) Analysis and Visualization of Tree Space. ed. by F. Anderson. Systematic Biology 54, 471–482. https://doi.org/10.1080/10635150590946961

Irisarri, I., Singh, P., Koblmüller, S., et al. (2018) Phylogenomics uncovers early hybridization and adaptive loci shaping the radiation of Lake Tanganyika cichlid fishes. Nature Communications 9. https://doi.org/10.1038/s41467-018-05479-9

Jacobs, A. & Therkildsen, N.O. (2019) Excavating ghost footprints and tangled trees from modern genomes. Molecular Ecology 28, 3287–3290. https://doi.org/10.1111/mec.15141

Joly, S., McLenachan, P.A. & Lockhart, P.J. (2009) A Statistical Approach for Distinguishing Hybridization and Incomplete Lineage Sorting. The American Naturalist 174, E54–E70. https://doi.org/10.1086/600082

Jones, M.R., Mills, L.S., Alves, P.C., Callahan, C.M., Alves, J.M., Lafferty, D.J.R., Jiggins, F.M., Jensen, J.D., Melo-Ferreira, J. & Good, J.M. (2018) Adaptive introgression underlies polymorphic seasonal camouflage in snowshoe hares. Science 360, 1355–1358. https://doi.org/10.1126/science.aar5273

Kendall, M. & Colijn, C. (2016) A tree metric using structure and length to capture distinct phylogenetic signals. Molecular Biology and Evolution 33, 2735–2743. https://doi.org/10.1093/molbev/msw124

Kim, B.-M., Amores, A., Kang, S., et al. (2019) Antarctic blackfin icefish genome reveals adaptations to extreme environments. Nature Ecology & Evolution 3, 469–478. https://doi.org/10.1038/s41559-019-0812-7

Kirkpatrick, M. (2010) How and Why Chromosome Inversions Evolve. PLOS Biology 8, e1000501. https://doi.org/10.1371/journal.pbio.1000501

Kirkpatrick, M. & Barton, N. (2006) Chromosome Inversions, Local Adaptation and Speciation. Genetics 173, 419–434.

Kryazhimskiy, S. & Plotkin, J.B. (2008) The Population Genetics of dN/dS. PLOS Genetics 4, e1000304. https://doi.org/10.1371/journal.pgen.1000304

Kuhlwilm, M., Han, S., Sousa, V.C., Excoffier, L. & Marques-Bonet, T. (2019) Ancient admixture from an extinct ape lineage into bonobos. Nature Ecology & Evolution 3, 957–965. https://doi.org/10.1038/s41559-019-0881-7

Lamichhaney, S., Berglund, J., Almén, M.S., et al. (2015) Evolution of Darwin’s finches and their beaks revealed by genome sequencing. Nature 518, nature14181. https://doi.org/10.1038/nature14181

Lehmann, R., Lightfoot, D.J., Schunter, C., et al. (2018) Finding Nemo’s Genes: A chromosome-scale reference assembly of the genome of the orange clownfish Amphiprion percula. Molecular Ecology Resources 19, 570–585. https://doi.org/10.1101/278267

Li, H. & Durbin, R. (2009) Fast and accurate short read alignment with Burrows–Wheeler transform. Bioinformatics 25, 1754–1760. https://doi.org/10.1093/bioinformatics/btp324

Link, V., Kousathanas, A., Veeramah, K., Sell, C., Scheu, A. & Wegmann, D. (2017) ATLAS: Analysis Tools for Low-depth and Ancient Samples. bioRxiv, 105346. https://doi.org/10.1101/105346

Litsios, G. & Salamin, N. (2014) Hybridisation and diversification in the adaptive radiation of clownfishes. BMC evolutionary biology 14, 245.

Litsios, G., Sims, C.A., Wüest, R.O., Pearman, P.B., Zimmermann, N.E. & Salamin, N. (2012) Mutualism with sea anemones triggered the adaptive radiation of clownfishes. BMC evolutionary biology 12, 212.

Lopes, F., Oliveira, L.R., Kessler, A., et al. (2021) Phylogenomic Discordance in the Eared Seals is best explained by Incomplete Lineage Sorting following Explosive Radiation in the Southern Hemisphere. ed. by J. Esselstyn. Systematic Biology 70, 786–802. https://doi.org/10.1093/sysbio/syaa099

Lowry, D.B. & Willis, J.H. (2010) A Widespread Chromosomal Inversion Polymorphism Contributes to a Major Life-History Transition, Local Adaptation, and Reproductive Isolation. PLoS Biology 8.

Malinsky, M., Matschiner, M. & Svardal, H. (2019) Dsuite - fast D-statistics and related admixture evidence from VCF files. Genomics. https://doi.org/10.1101/634477

Malinsky, M., Svardal, H., Tyers, A.M., Miska, E.A., Genner, M.J., Turner, G.F. & Durbin, R. (2018) Whole-genome sequences of Malawi cichlids reveal multiple radiations interconnected by gene flow. Nature Ecology & Evolution 2, 1940–1955. https://doi.org/10.1038/s41559-018-0717-x

Mallet, J. (2007) Hybrid speciation. Nature 446, 279–283.

Mallet, J., Beltrán, M., Neukirchen, W. & Linares, M. (2007) Natural hybridization in heliconiine butterflies: the species boundary as a continuum. BMC Evolutionary Biology 7, 28. https://doi.org/10.1186/1471-2148-7-28

Marcionetti, A., Rossier, V., Roux, N., Salis, P., Laudet, V. & Salamin, N. (2019) Insights into the Genomics of Clownfish Adaptive Radiation: Genetic Basis of the Mutualism with Sea Anemones. Genome Biology and Evolution 11, 869–882. https://doi.org/10.1093/gbe/evz042

Marcionetti, A. & Salamin, N. (2022) Insights into the genomics of clownfish adaptive radiation: the genomic substrate of the diversification. bioRxiv, 2022.05.12.491701. https://doi.org/10.1101/2022.05.12.491701

Marin, J., Achaz, G., Crombach, A. & Lambert, A. (2020) The genomic view of diversification. Journal of Evolutionary Biology. https://doi.org/10.1111/jeb.13677

Marques, D.A., Meier, J.I. & Seehausen, O. (2019) A Combinatorial View on Speciation and Adaptive Radiation. Trends in Ecology & Evolution. https://doi.org/10.1016/j.tree.2019.02.008

Martin R. Smith (2020) TreeDist: Distances Between Phylogenetic Trees. Zenodo. https://doi.org/10.5281/zenodo.3941734

Martin, S.H., Davey, J.W. & Jiggins, C.D. (2015) Evaluating the Use of ABBA–BABA Statistics to Locate Introgressed Loci. Molecular Biology and Evolution 32, 244–257. https://doi.org/10.1093/molbev/msu269

Meier, J.I., Marques, D.A., Mwaiko, S., Wagner, C.E., Excoffier, L. & Seehausen, O. (2017) Ancient hybridization fuels rapid cichlid fish adaptive radiations. Nature Communications 8. https://doi.org/10.1038/ncomms14363

Meier, J.I., Marques, D.A., Wagner, C.E., Excoffier, L. & Seehausen, O. (2018) Genomics of Parallel Ecological Speciation in Lake Victoria Cichlids. Molecular Biology and Evolution 35, 1489–1506. https://doi.org/10.1093/molbev/msy051

Meleshko, O., Martin, M.D., Korneliussen, T.S., et al. (2021) Extensive Genome-Wide Phylogenetic Discordance Is Due to Incomplete Lineage Sorting and Not Ongoing Introgression in a Rapidly Radiated Bryophyte Genus. ed. by J. de Meaux. Molecular Biology and Evolution 38, 2750–2766. https://doi.org/10.1093/molbev/msab063

Minh, B.Q., Schmidt, H.A., Chernomor, O., Schrempf, D., Woodhams, M.D., von Haeseler, A. & Lanfear, R. (2020) IQ-TREE 2: New Models and Efficient Methods for Phylogenetic Inference in the Genomic Era. Molecular Biology and Evolution 37, 1530–1534. https://doi.org/10.1093/molbev/msaa015

Natri, H.M., Merilä, J. & Shikano, T. (2019) The evolution of sex determination associated with a chromosomal inversion. Nature Communications 10, 145. https://doi.org/10.1038/s41467-018-08014-y

Ottenburghs, J. (2020) Ghost Introgression: Spooky Gene Flow in the Distant Past. BioEssays, 2000012. https://doi.org/10.1002/bies.202000012

Paradis, E. & Schliep, K. (2019) ape 5.0: an environment for modern phylogenetics and evolutionary analyses in R. Bioinformatics 35, 526–528. https://doi.org/10.1093/bioinformatics/bty633

Pardo-Diaz, C., Salazar, C., Baxter, S.W., Merot, C., Figueiredo-Ready, W., Joron, M., McMillan, W.O. & Jiggins, C.D. (2012) Adaptive Introgression across Species Boundaries in Heliconius Butterflies. PLOS Genetics 8, e1002752. https://doi.org/10.1371/journal.pgen.1002752

Payseur, B.A. & Rieseberg, L.H. (2016) A genomic perspective on hybridization and speciation. Molecular Ecology 25, 2337–2360. https://doi.org/10.1111/mec.13557

Poelstra, J.W., Richards, E.J. & Martin, C.H. (2018) Speciation in sympatry with ongoing secondary gene flow and a potential olfactory trigger in a radiation of Cameroon cichlids. Molecular Ecology 27, 4270– 4288. https://doi.org/10.1111/mec.14784

Rafati, N., Blanco-Aguiar, J.A., Rubin, C.J., et al. (2018) A genomic map of clinal variation across the European rabbit hybrid zone. Molecular Ecology 27, 1457–1478. https://doi.org/10.1111/mec.14494

Runemark, A., Vallejo-Marin, M. & Meier, J.I. (2019) Eukaryote hybrid genomes. PLOS Genetics 15, e1008404. https://doi.org/10.1371/journal.pgen.1008404

Salzburger, W. (2018) Understanding explosive diversification through cichlid fish genomics. Nature Reviews Genetics 19, 705–717. https://doi.org/10.1038/s41576-018-0043-9

Sankararaman, S., Mallick, S., Patterson, N. & Reich, D. (2016) The Combined Landscape of Denisovan and Neanderthal Ancestry in Present-Day Humans. Current Biology 26, 1241–1247. https://doi.org/10.1016/j.cub.2016.03.037

Schluter, D. (2000) The Ecology of Adaptive Radiation. Oxford Series in Ecology and Evolution.,. Oxford University Press.

Schumer, M., Cui, R., Powell, D.L., Rosenthal, G.G. & Andolfatto, P. (2016) Ancient hybridization and genomic stabilization in a swordtail fish. Molecular Ecology 25, 2661–2679. https://doi.org/10.1111/mec.13602

Seehausen, O. (2004) Hybridization and adaptive radiation. Trends in Ecology & Evolution 19, 198–207. https://doi.org/10.1016/j.tree.2004.01.003

Sievers, F. & Higgins, D.G. (2014) Clustal Omega, accurate alignment of very large numbers of sequences. Methods in Molecular Biology (Clifton, N.J.) 1079, 105–116. https://doi.org/10.1007/978-1-62703-646-7_6

Smith, S.A., Brown, J.W. & Walker, J.F. (2018) So many genes, so little time: A practical approach to divergence-time estimation in the genomic era. PLOS ONE 13, e0197433. https://doi.org/10.1371/journal.pone.0197433

Stamatakis, A. (2014) RAxML version 8: a tool for phylogenetic analysis and post-analysis of large phylogenies. Bioinformatics 30, 1312–1313. https://doi.org/10.1093/bioinformatics/btu033

Stevison, L.S., Hoehn, K.B. & Noor, M.A.F. (2011) Effects of inversions on within- and between-species recombination and divergence. Genome Biology and Evolution 3, 830–841. https://doi.org/10.1093/gbe/evr081

Svardal, H., Quah, F.X., Malinsky, M., Ngatunga, B.P., Miska, E.A., Salzburger, W., Genner, M.J., Turner, G.F. & Durbin, R. (2020) Ancestral Hybridization Facilitated Species Diversification in the Lake Malawi Cichlid Fish Adaptive Radiation. ed. by R. Nielsen. Molecular Biology and Evolution 37, 1100–1113. https://doi.org/10.1093/molbev/msz294

Wallbank, R.W.R., Baxter, S.W., Pardo-Diaz, C., et al. (2016) Evolutionary Novelty in a Butterfly Wing Pattern through Enhancer Shuffling. ed. by N.H. Barton. PLOS Biology 14, e1002353. https://doi.org/10.1371/journal.pbio.1002353

Wang, K., Lenstra, J.A., Liu, L., Hu, Q., Ma, T., Qiu, Q. & Liu, J. (2018) Incomplete lineage sorting rather than hybridization explains the inconsistent phylogeny of the wisent. Communications Biology 1, 1–9. https://doi.org/10.1038/s42003-018-0176-6

Weir, J.T. & Price, M. (2011) Andean uplift promotes lowland speciation through vicariance and dispersal in Dendrocincla woodcreepers. Molecular Ecology 20, 4550–4563. https://doi.org/10.1111/j.1365-294X.2011.05294.x

Whitney, K.D., Randel, R.A. & Rieseberg, L.H. (2010) Adaptive introgression of abiotic tolerance traits in the sunflower Helianthus annuus. New Phytologist 187, 230–239. https://doi.org/10.1111/j.1469-8137.2010.03234.x

Yu, Y., Barnett, R.M. & Nakhleh, L. (2013) Parsimonious Inference of Hybridization in the Presence of Incomplete Lineage Sorting. Systematic Biology 62, 738–751. https://doi.org/10.1093/sysbio/syt037

Zhang, C., Rabiee, M., Sayyari, E. & Mirarab, S. (2018) ASTRAL-III: polynomial time species tree reconstruction from partially resolved gene trees. BMC Bioinformatics 19, 153. https://doi.org/10.1186/s12859-018-2129-y

