## Supplementary figures and images for "Extensive hybridisation throughout clownfishes evolutionary history"

### Supplemental Figure 1

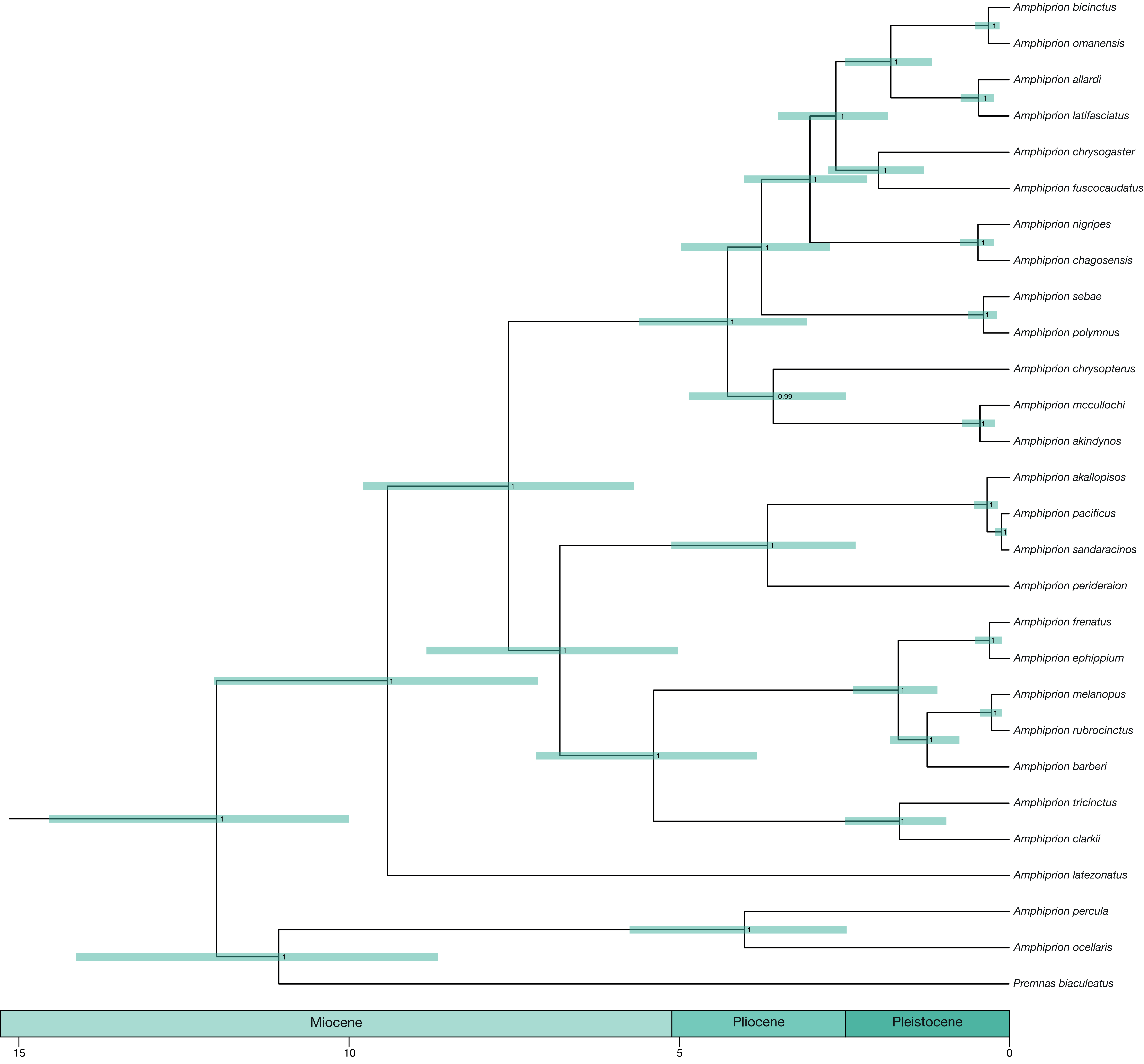

### Supplemental Figure 3

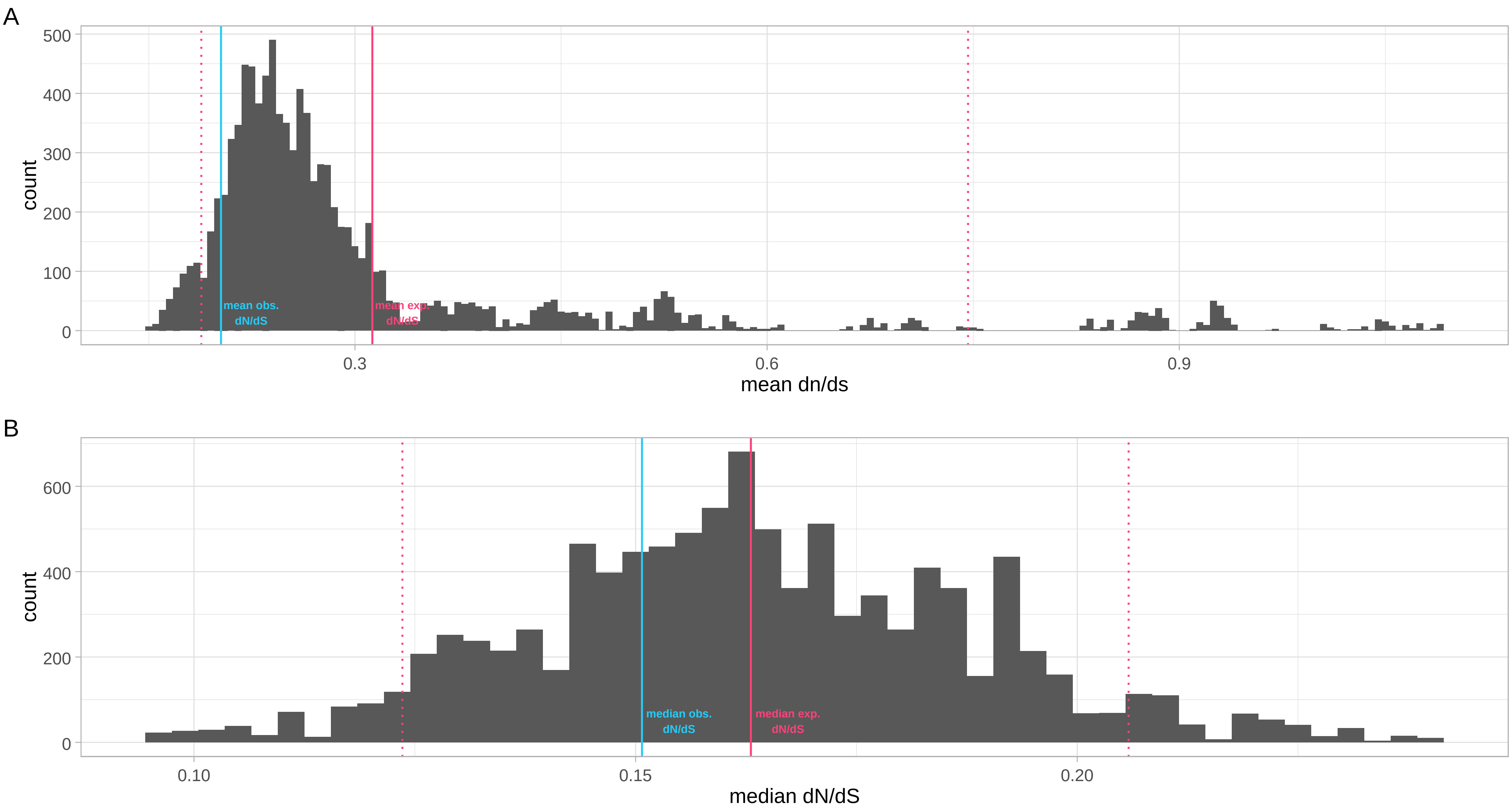
