## Supplemental Figure 2 for "Extensive hybridisation throughout clownfishes evolutionary history"

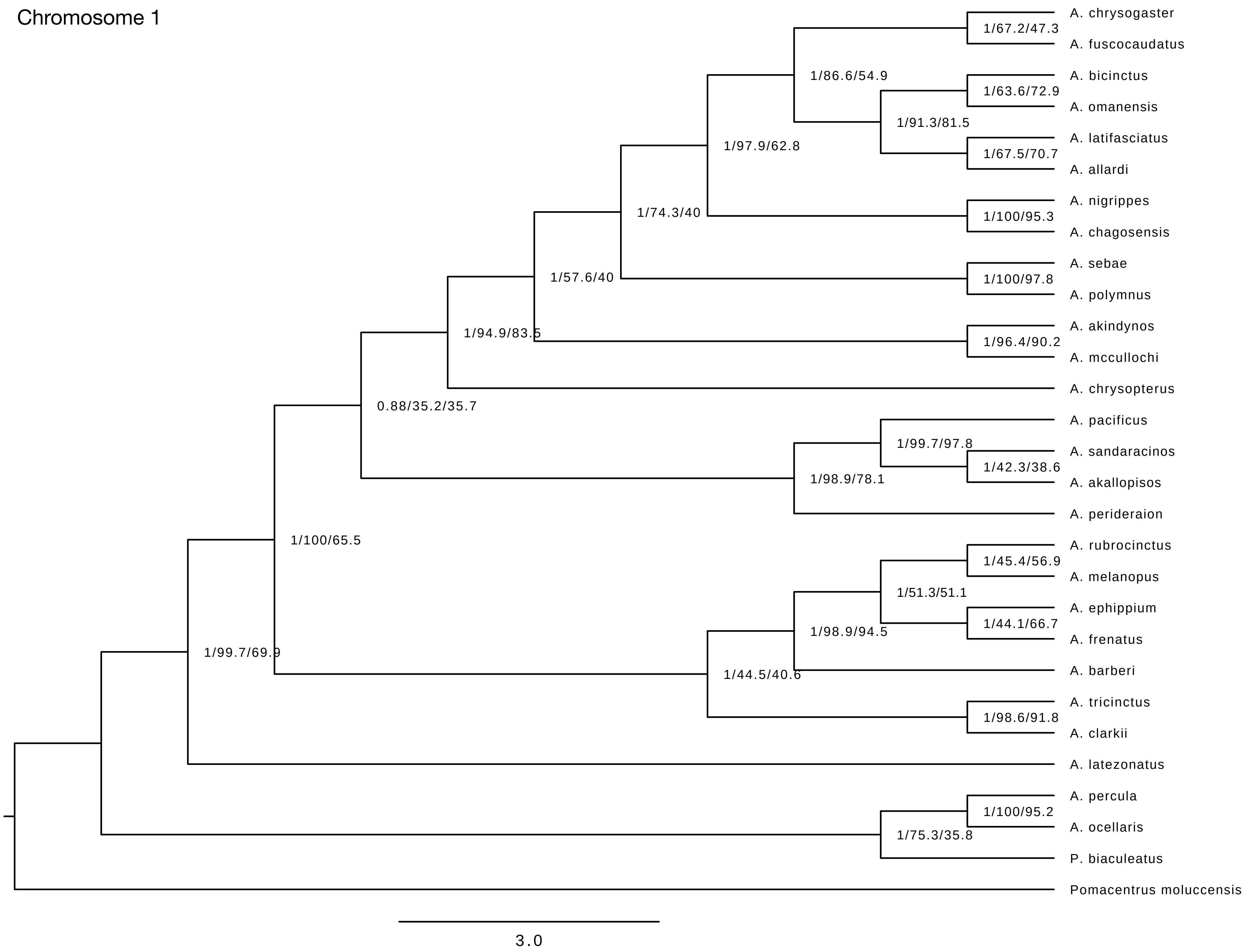
Chromosome 1
A. chrysogaster
A. fuscocaudatus
A. bicinctus
A. omanensis
A. latifasciatus
A. allardi
A. nigrippes
A. chagosensis
A. sebae
A. polymnus
A. akindynos
A. mccullochi
A. chrysopterus
A. pacificus
A. sandaracinos
A. akallopisos
A. perideraion
A. rubrocinctus
A. melanopus
A. ephippium
A. frenatus
A. barberi
A. tricinctus
A. clarkii
A. latezonatus
A. percula
A. ocellaris
P. biaculeatus
Pomacentrus moluccensis
1/67.2/47.3
1/86.6/54.9
1/63.6/72.9
1/91.3/81.5
1/67.5/70.7
1/97.9/62.8
1/74.3/40
1/100/95.3
1/57.6/40
1/100/97.8
1/94.9/83.5
1/96.4/90.2
0.88/35.2/35.7
1/99.7/97.8
1/42.3/38.6
1/98.9/78.1
1/100/65.5
1/45.4/56.9
1/51.3/51.1
1/44.1/66.7
1/98.9/94.5
1/99.7/69.9
1/44.5/40.6
1/98.6/91.8
1/100/95.2
1/75.3/35.8
3.0

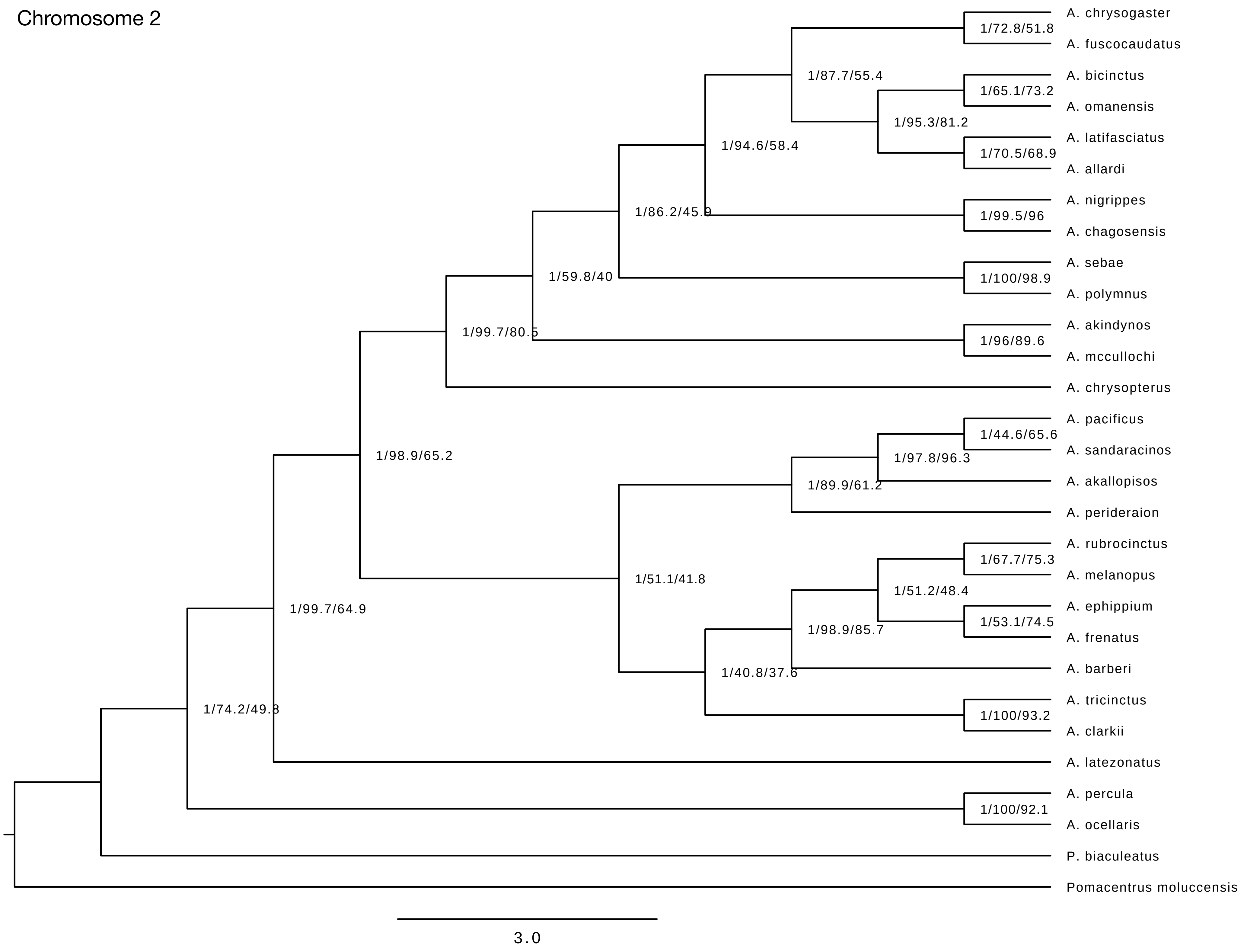
Chromosome 2
A. chrysogaster
A. fuscocaudatus
A. bicinctus
A. omanensis
A. latifasciatus
A. allardi
A. nigrippes
A. chagosensis
A. sebae
A. polymnus
A. akindynos
A. mccullochi
A. chrysopterus
A. pacificus
A. sandaracinos
A. akallopisos
A. perideraion
A. rubrocinctus
A. melanopus
A. ephippium
A. frenatus
A. barberi
A. tricinctus
A. clarkii
A. latezonatus
A. percula
A. ocellaris
P. biaculeatus
Pomacentrus moluccensis
1/72.8/51.8
1/87.7/55.4
1/65.1/73.2
1/95.3/81.2
1/94.6/58.4
1/70.5/68.9
1/99.5/96
1/86.2/45.9
1/59.8/40
1/100/98.9
1/99.7/80.5
1/96/89.6
1/98.9/65.2
1/44.6/65.6
1/97.8/96.3
1/89.9/61.2
1/67.7/75.3
1/51.1/41.8
1/51.2/48.4
1/99.7/64.9
1/53.1/74.5
1/98.9/85.7
1/40.8/37.6
1/74.2/49.3
1/100/93.2
1/100/92.1
3.0

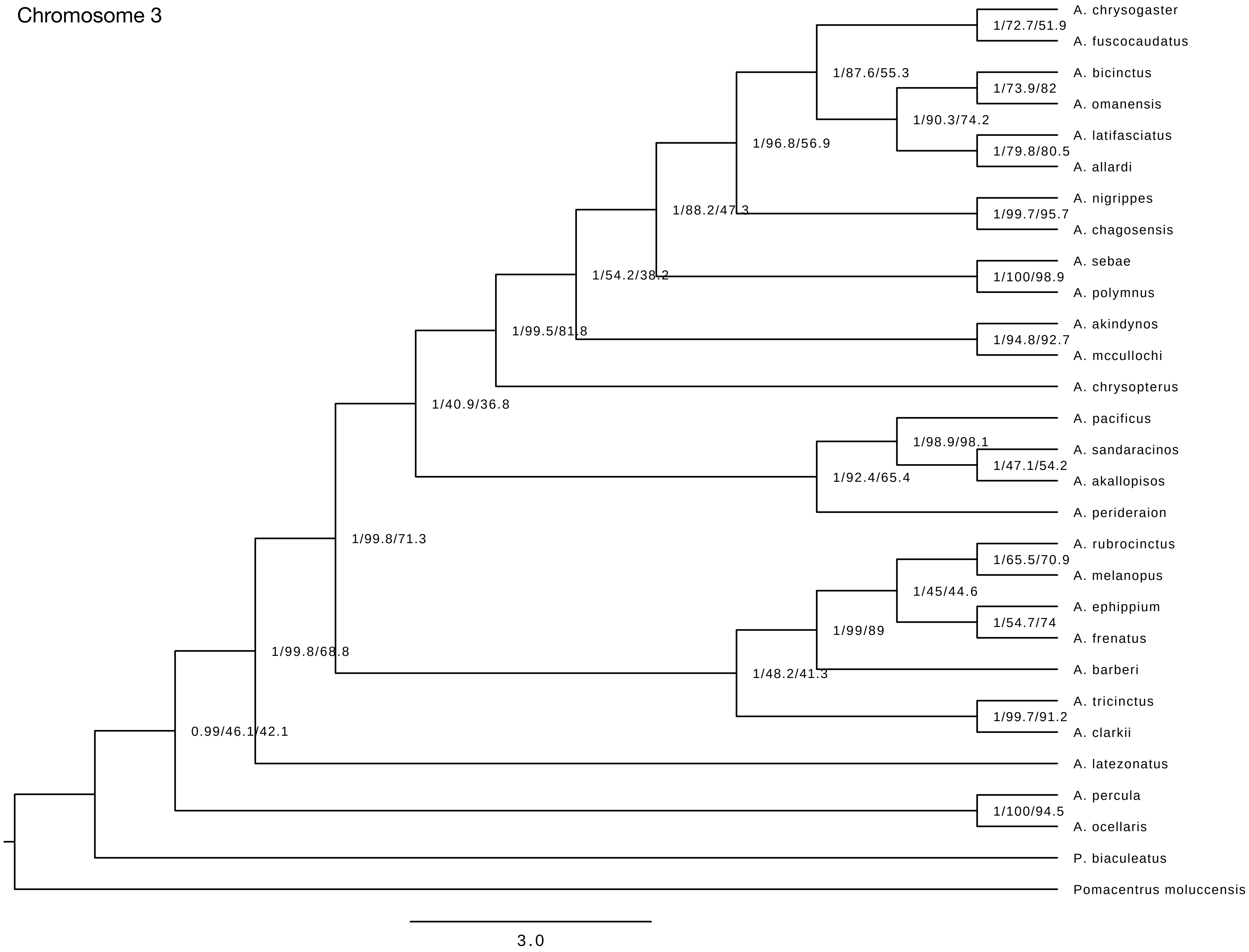
Chromosome 3
A. chrysogaster
1/72.7/51.9
A. fuscocaudatus
1/87.6/55.3
A. bicinctus
1/73.9/82
A. omanensis
1/90.3/74.2
A. latifasciatus
1/96.8/56.9
1/79.8/80.5
A. allardi
A. nigrippes
1/88.2/47.3
1/99.7/95.7
A. chagosensis
A. sebae
1/54.2/38.2
1/100/98.9
A. polymnus
A. akindynos
1/99.5/81.8
1/94.8/92.7
A. mccullochi
A. chrysopterus
1/40.9/36.8
A. pacificus
1/98.9/98.1
A. sandaracinos
1/47.1/54.2
1/92.4/65.4
A. akallopisos
A. perideraion
1/99.8/71.3
A. rubrocinctus
1/65.5/70.9
A. melanopus
1/45/44.6
A. ephippium
1/54.7/74
1/99/89
A. frenatus
1/99.8/68.8
A. barberi
1/48.2/41.3
A. tricinctus
1/99.7/91.2
0.99/46.1/42.1
A. clarkii
A. latezonatus
A. percula
1/100/94.5
A. ocellaris
P. biaculeatus
Pomacentrus moluccensis
3.0

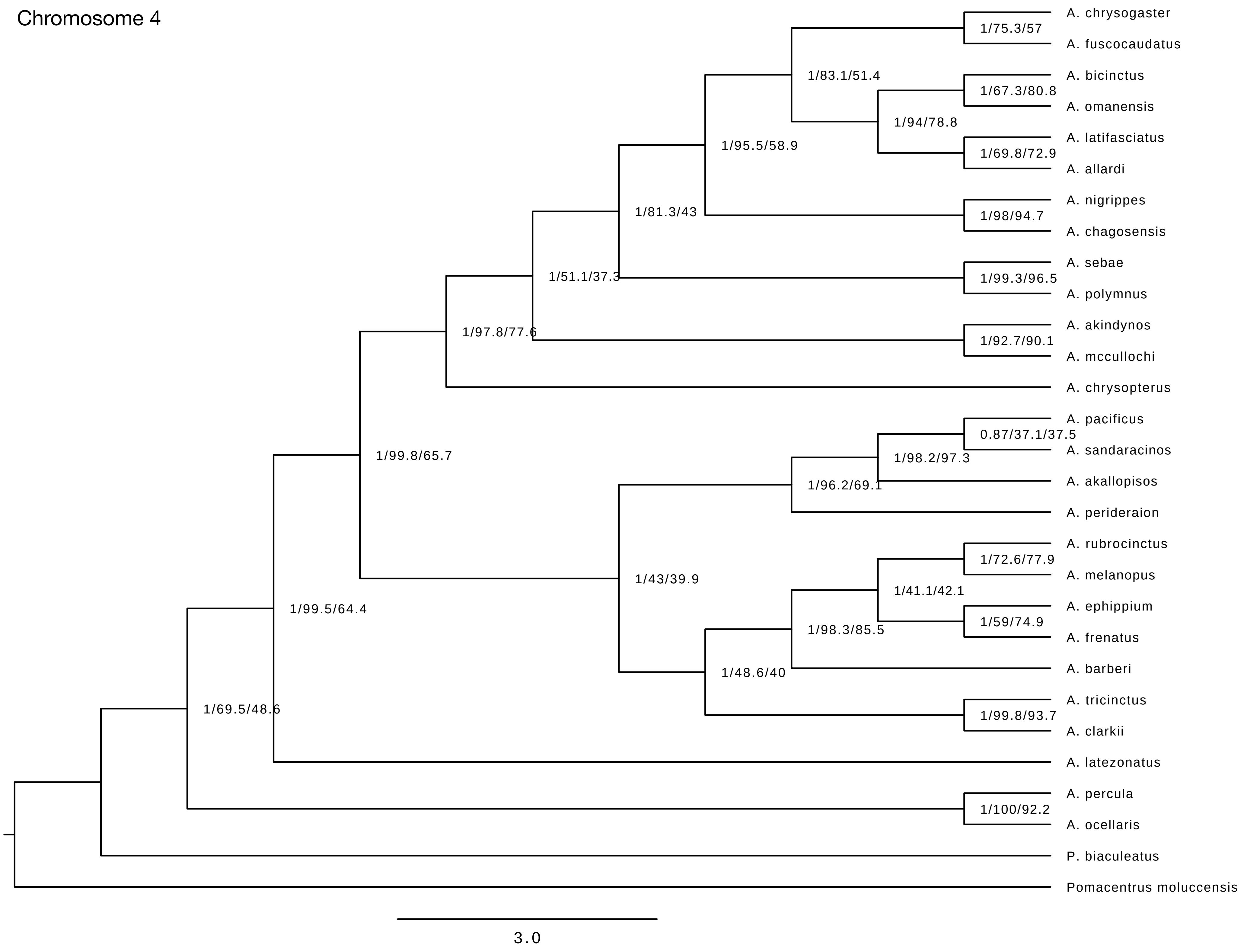
Chromosome 4
A. chrysogaster
A. fuscocaudatus
A. bicinctus
A. omanensis
A. latifasciatus
A. allardi
A. nigrippes
A. chagosensis
A. sebae
A. polymnus
A. akindynos
A. mccullochi
A. chrysopterus
A. pacificus
A. sandaracinos
A. akallopisos
A. perideraion
A. rubrocinctus
A. melanopus
A. ephippium
A. frenatus
A. barberi
A. tricinctus
A. clarkii
A. latezonatus
A. percula
A. ocellaris
P. biaculeatus
Pomacentrus moluccensis
1/75.3/57
1/83.1/51.4
1/67.3/80.8
1/94/78.8
1/69.8/72.9
1/95.5/58.9
1/98/94.7
1/81.3/43
1/99.3/96.5
1/51.1/37.3
1/92.7/90.1
1/97.8/77.6
0.87/37.1/37.5
1/98.2/97.3
1/96.2/69.1
1/99.8/65.7
1/72.6/77.9
1/41.1/42.1
1/59/74.9
1/98.3/85.5
1/43/39.9
1/48.6/40
1/99.8/93.7
1/99.5/64.4
1/69.5/48.6
1/100/92.2
3.0

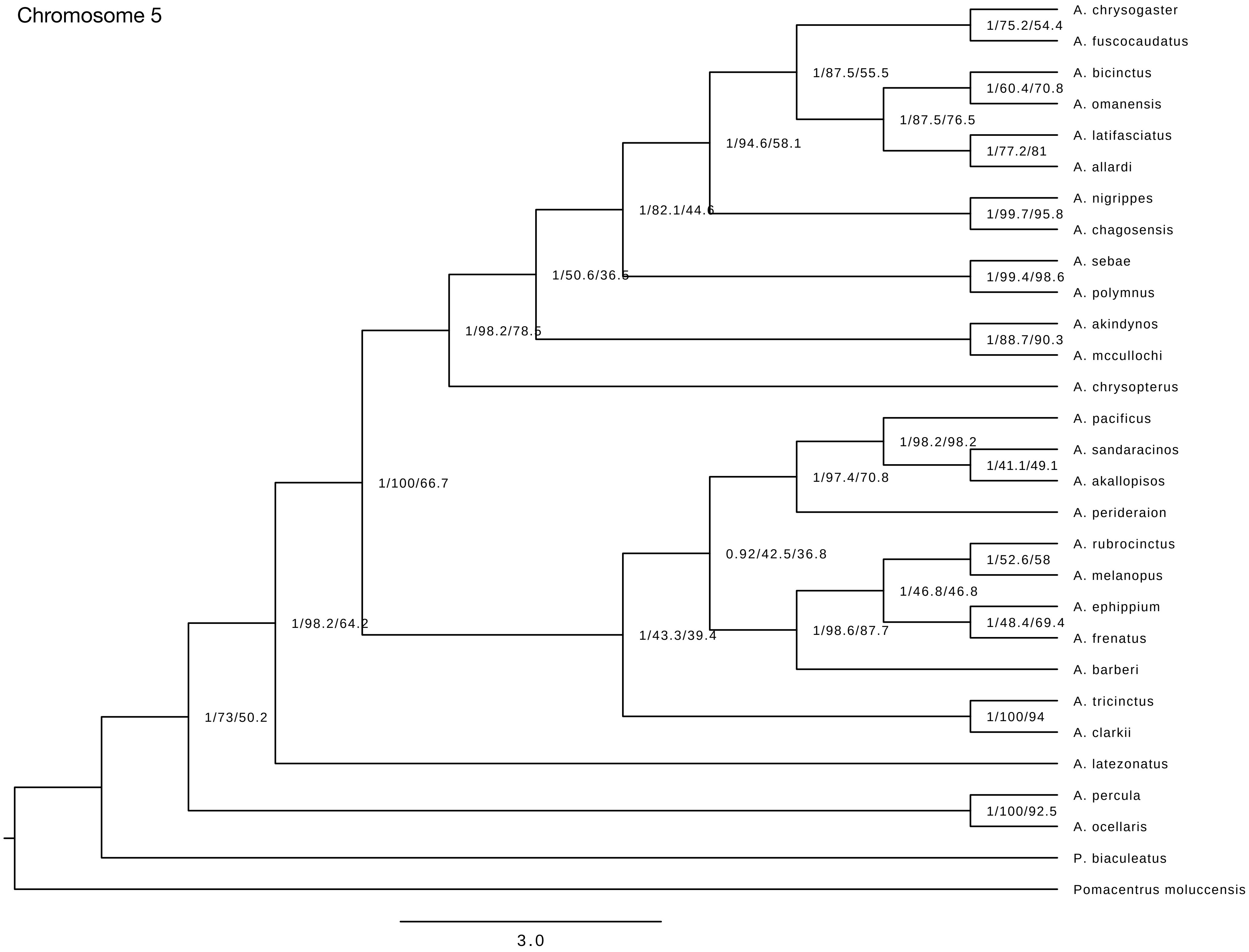
Chromosome 5
A. chrysogaster
A. fuscocaudatus
1/75.2/54.4
A. bicinctus
A. omanensis
1/60.4/70.8
A. latifasciatus
A. allardi
1/77.2/81
1/87.5/76.5
1/87.5/55.5
1/94.6/58.1
A. nigrippes
A. chagosensis
1/99.7/95.8
1/82.1/44.6
A. sebae
A. polymnus
1/99.4/98.6
1/50.6/36.5
A. akindynos
A. mccullochi
1/88.7/90.3
1/98.2/78.5
A. chrysopterus
1/100/66.7
A. pacificus
A. sandaracinos
A. akallopisos
1/41.1/49.1
1/98.2/98.2
1/97.4/70.8
A. perideraion
0.92/42.5/36.8
A. rubrocinctus
A. melanopus
1/52.6/58
A. ephippium
A. frenatus
1/48.4/69.4
1/46.8/46.8
1/98.6/87.7
A. barberi
1/43.3/39.4
A. tricinctus
A. clarkii
1/100/94
1/98.2/64.2
A. latezonatus
1/73/50.2
A. percula
A. ocellaris
1/100/92.5
P. biaculeatus
Pomacentrus moluccensis
3.0

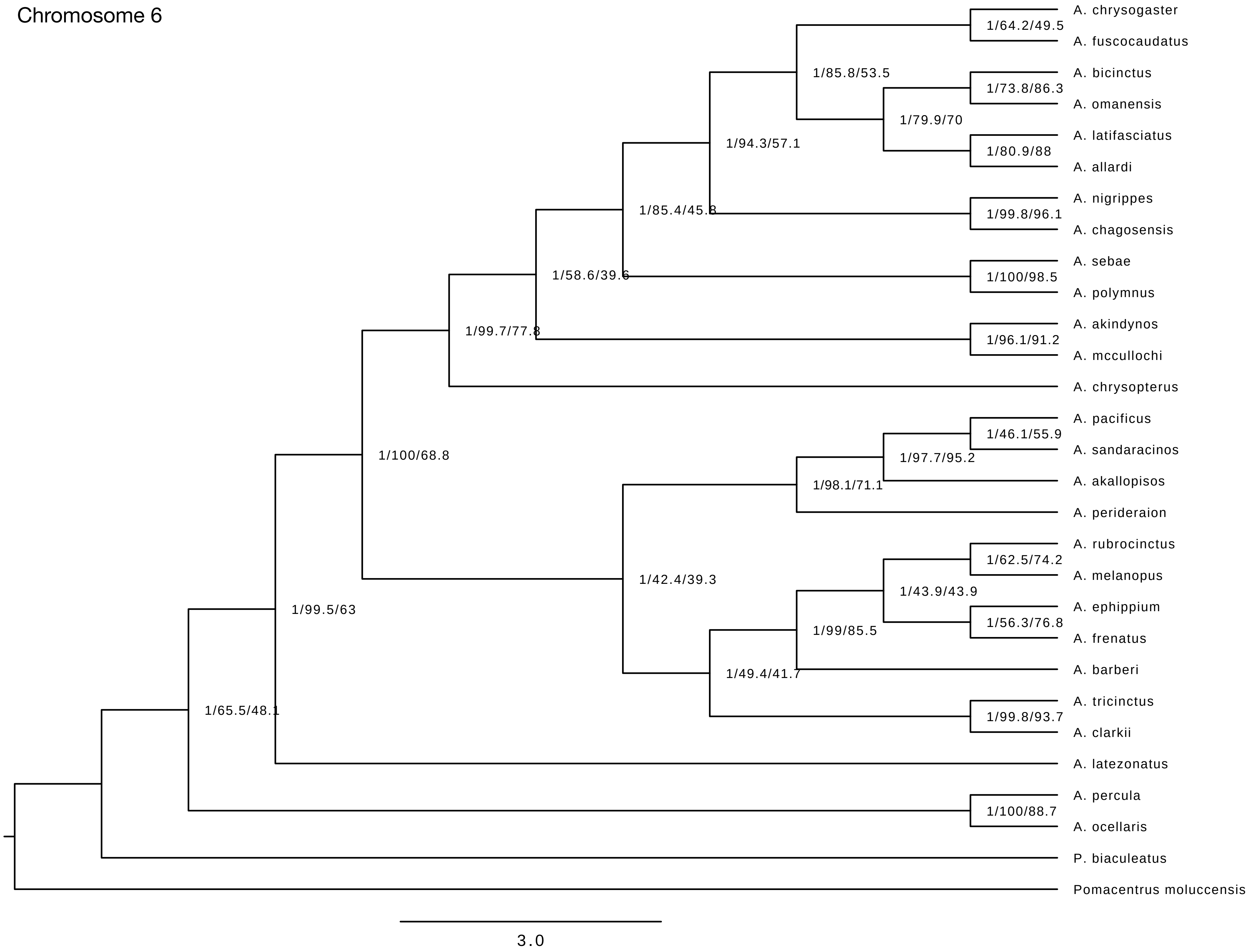
Chromosome 6
A. chrysogaster
A. fuscocaudatus
A. bicinctus
A. omanensis
A. latifasciatus
A. allardi
A. nigrippes
A. chagosensis
A. sebae
A. polymnus
A. akindynos
A. mccullochi
A. chrysopterus
A. pacificus
A. sandaracinos
A. akallopisos
A. perideraion
A. rubrocinctus
A. melanopus
A. ephippium
A. frenatus
A. barberi
A. tricinctus
A. clarkii
A. latezonatus
A. percula
A. ocellaris
P. biaculeatus
Pomacentrus moluccensis
1/64.2/49.5
1/85.8/53.5
1/73.8/86.3
1/79.9/70
1/80.9/88
1/94.3/57.1
1/85.4/45.8
1/99.8/96.1
1/58.6/39.6
1/100/98.5
1/99.7/77.8
1/96.1/91.2
1/100/68.8
1/46.1/55.9
1/97.7/95.2
1/98.1/71.1
1/62.5/74.2
1/43.9/43.9
1/42.4/39.3
1/56.3/76.8
1/99.5/63
1/99/85.5
1/49.4/41.7
1/65.5/48.1
1/99.8/93.7
1/100/88.7
3.0

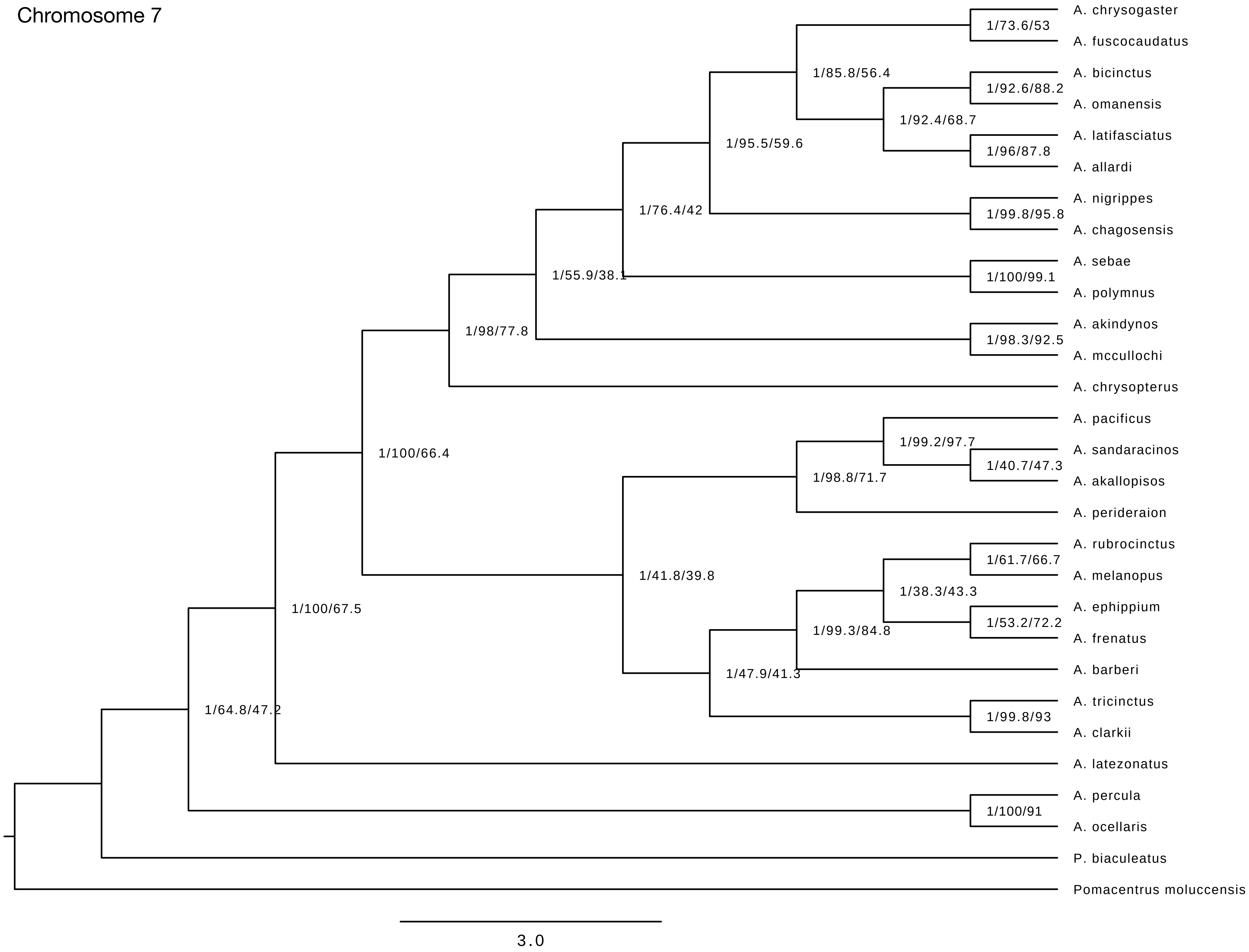
Chromosome 7
A. chrysogaster
A. fuscocaudatus
A. bicinctus
A. omanensis
A. latifasciatus
A. allardi
A. nigrippes
A. chagosensis
A. sebae
A. polymnus
A. akindynos
A. mccullochi
A. chrysopterus
A. pacificus
A. sandaracinos
A. akallopisos
A. perideraion
A. rubrocinctus
A. melanopus
A. ephippium
A. frenatus
A. barberi
A. tricinctus
A. clarkii
A. latezonatus
A. percula
A. ocellaris
P. biaculeatus
Pomacentrus moluccensis
1/73.6/53
1/85.8/56.4
1/92.6/88.2
1/92.4/68.7
1/96/87.8
1/95.5/59.6
1/76.4/42
1/99.8/95.8
1/55.9/38.1
1/100/99.1
1/98/77.8
1/98.3/92.5
1/100/66.4
1/99.2/97.7
1/40.7/47.3
1/98.8/71.7
1/41.8/39.8
1/61.7/66.7
1/38.3/43.3
1/100/67.5
1/53.2/72.2
1/99.3/84.8
1/47.9/41.3
1/64.8/47.2
1/99.8/93
1/100/91
3.0

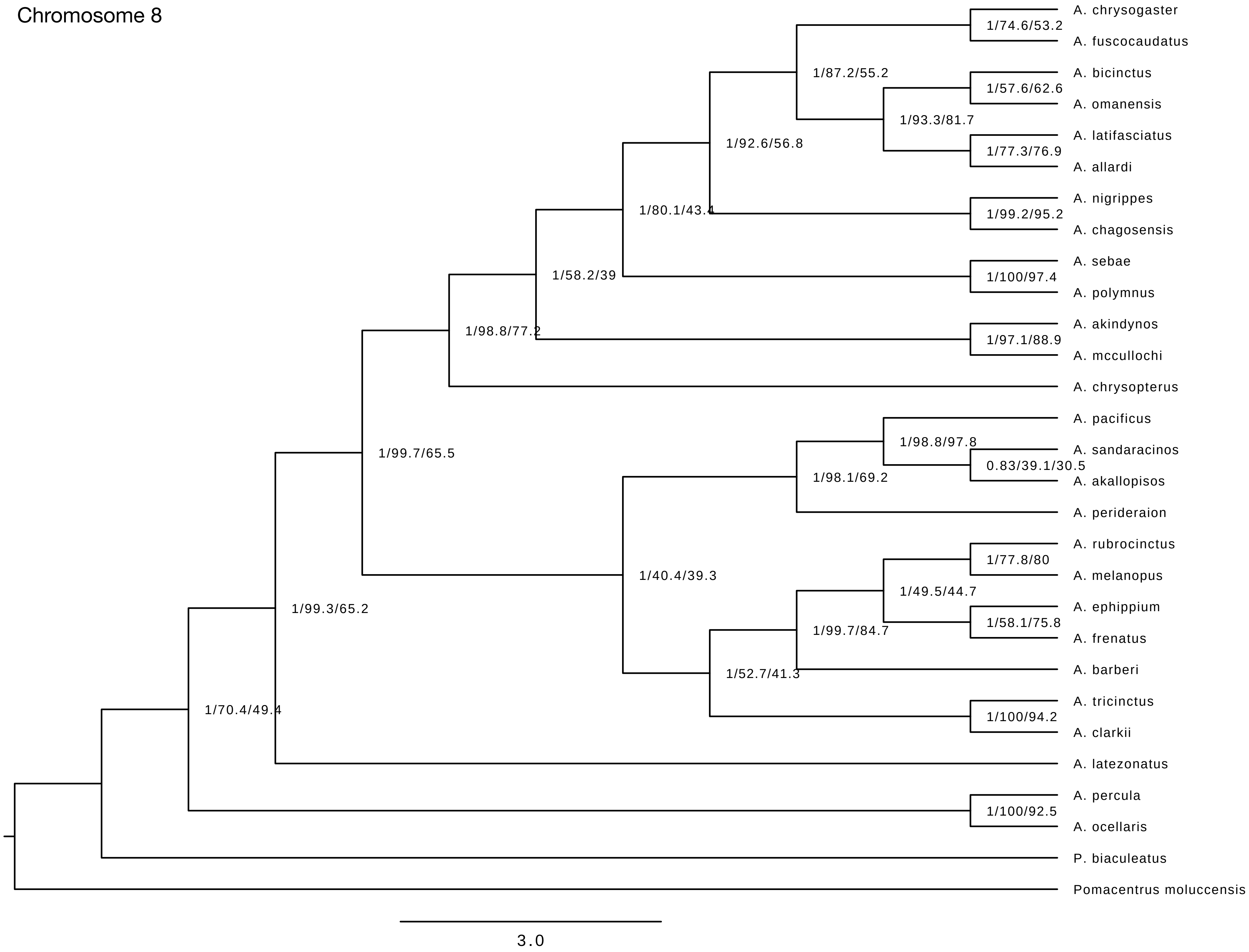
Chromosome 8
A. chrysogaster
A. fuscocaudatus
A. bicinctus
A. omanensis
A. latifasciatus
A. allardi
A. nigrippes
A. chagosensis
A. sebae
A. polymnus
A. akindynos
A. mccullochi
A. chrysopterus
A. pacificus
A. sandaracinos
A. akallopisos
A. perideraion
A. rubrocinctus
A. melanopus
A. ephippium
A. frenatus
A. barberi
A. tricinctus
A. clarkii
A. latezonatus
A. percula
A. ocellaris
P. biaculeatus
Pomacentrus moluccensis
1/74.6/53.2
1/87.2/55.2
1/57.6/62.6
1/93.3/81.7
1/77.3/76.9
1/92.6/56.8
1/99.2/95.2
1/80.1/43.4
1/100/97.4
1/58.2/39
1/97.1/88.9
1/98.8/77.2
1/98.8/97.8
0.83/39.1/30.5
1/98.1/69.2
1/99.7/65.5
1/77.8/80
1/40.4/39.3
1/49.5/44.7
1/99.3/65.2
1/58.1/75.8
1/99.7/84.7
1/52.7/41.3
1/100/94.2
1/70.4/49.4
1/100/92.5
3.0

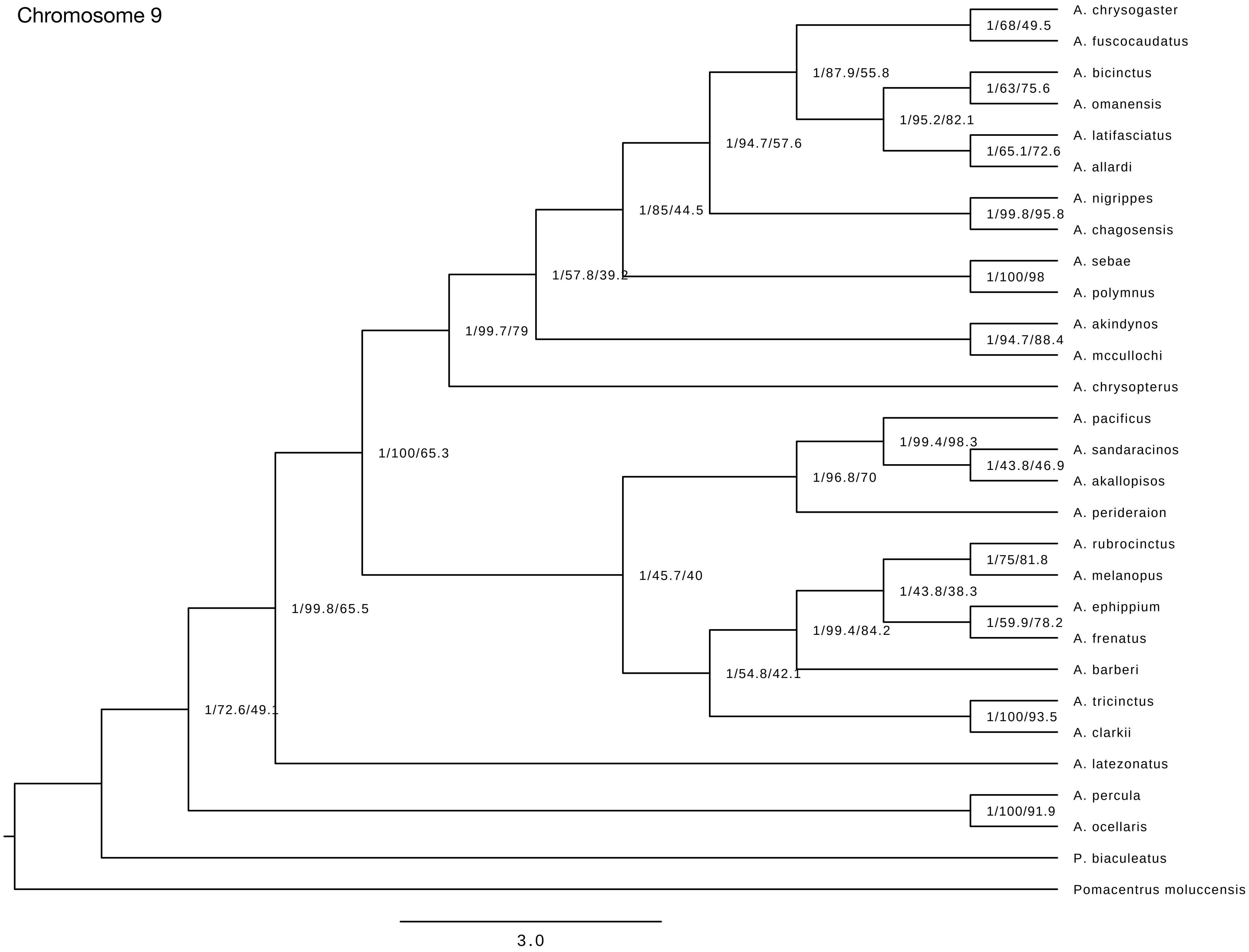
Chromosome 9
A. chrysogaster
A. fuscocaudatus
A. bicinctus
A. omanensis
A. latifasciatus
A. allardi
A. nigrippes
A. chagosensis
A. sebae
A. polymnus
A. akindynos
A. mccullochi
A. chrysopterus
A. pacificus
A. sandaracinos
A. akallopisos
A. perideraion
A. rubrocinctus
A. melanopus
A. ephippium
A. frenatus
A. barberi
A. tricinctus
A. clarkii
A. latezonatus
A. percula
A. ocellaris
P. biaculeatus
Pomacentrus moluccensis
1/68/49.5
1/87.9/55.8
1/63/75.6
1/95.2/82.1
1/65.1/72.6
1/94.7/57.6
1/85/44.5
1/99.8/95.8
1/57.8/39.2
1/100/98
1/99.7/79
1/94.7/88.4
1/100/65.3
1/99.4/98.3
1/43.8/46.9
1/96.8/70
1/75/81.8
1/45.7/40
1/43.8/38.3
1/99.8/65.5
1/59.9/78.2
1/99.4/84.2
1/54.8/42.1
1/72.6/49.1
1/100/93.5
1/100/91.9
3.0

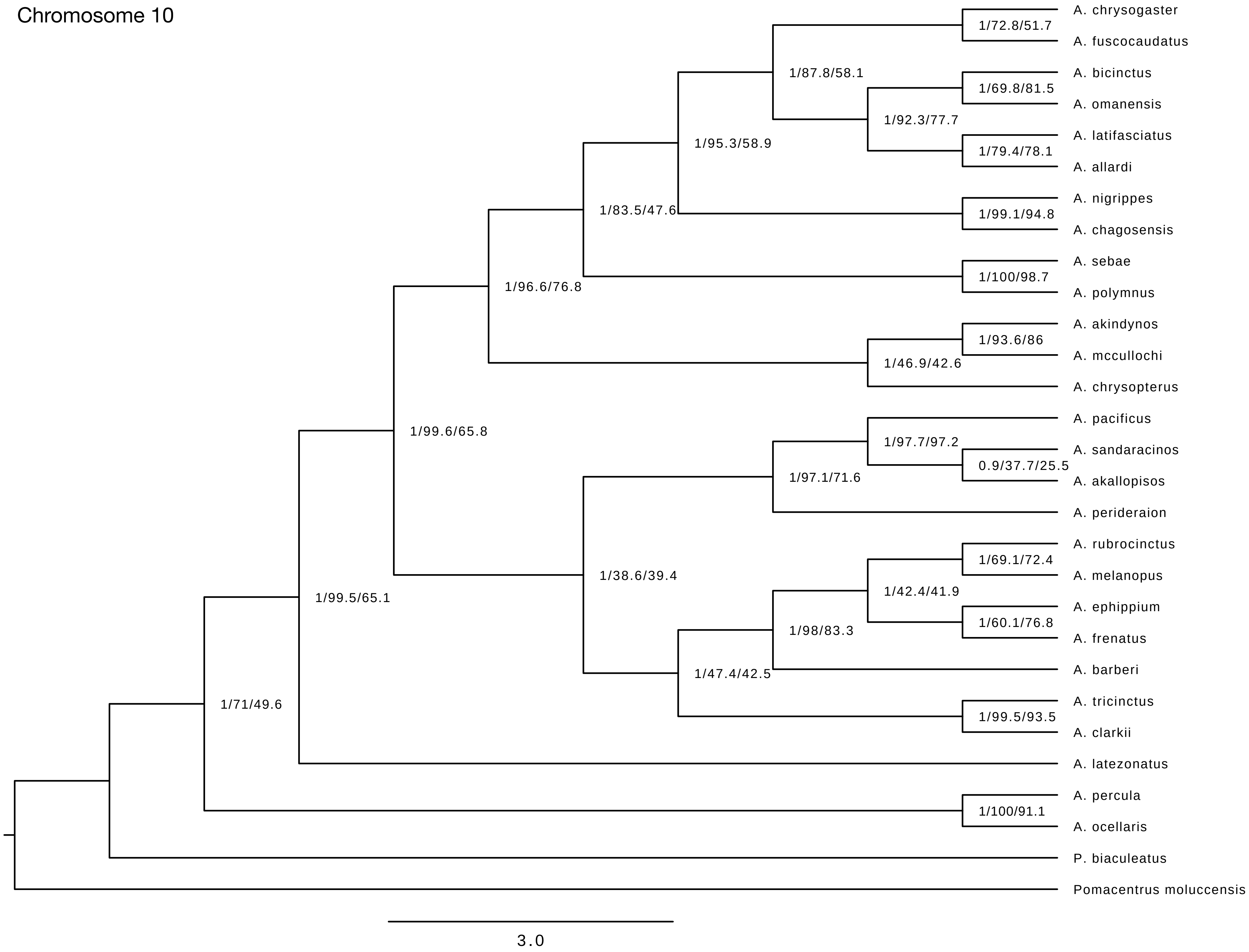

Chromosome 10
A. chrysogaster
A. fuscocaudatus
A. bicinctus
A. omanensis
A. latifasciatus
A. allardi
A. nigrippes
A. chagosensis
A. sebae
A. polymnus
A. akindynos
A. mccullochi
A. chrysopterus
A. pacificus
A. sandaracinos
A. akallopisos
A. perideraion
A. rubrocinctus
A. melanopus
A. ephippium
A. frenatus
A. barberi
A. tricinctus
A. clarkii
A. latezonatus
A. percula
A. ocellaris
P. biaculeatus
Pomacentrus moluccensis
1/72.8/51.7
1/87.8/58.1
1/69.8/81.5
1/92.3/77.7
1/95.3/58.9
1/79.4/78.1
1/83.5/47.6
1/99.1/94.8
1/100/98.7
1/96.6/76.8
1/93.6/86
1/46.9/42.6
1/99.6/65.8
1/97.7/97.2
0.9/37.7/25.5
1/97.1/71.6
1/69.1/72.4
1/38.6/39.4
1/42.4/41.9
1/99.5/65.1
1/98/83.3
1/60.1/76.8
1/47.4/42.5
1/71/49.6
1/99.5/93.5
1/100/91.1
3.0

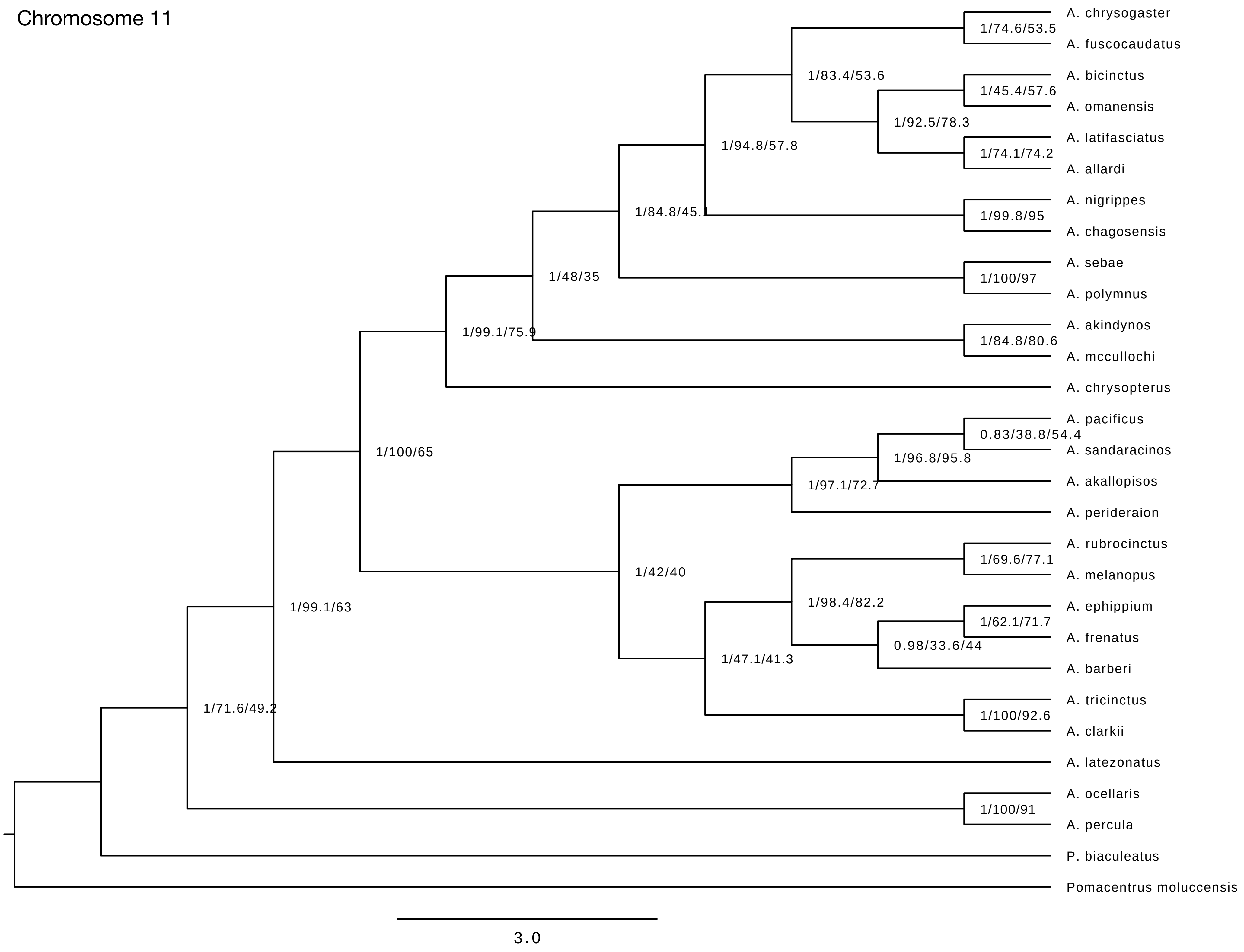
Chromosome 11
A. chrysogaster
A. fuscocaudatus
A. bicinctus
A. omanensis
A. latifasciatus
A. allardi
A. nigrippes
A. chagosensis
A. sebae
A. polymnus
A. akindynos
A. mccullochi
A. chrysopterus
A. pacificus
A. sandaracinos
A. akallopisos
A. perideraion
A. rubrocinctus
A. melanopus
A. ephippium
A. frenatus
A. barberi
A. tricinctus
A. clarkii
A. latezonatus
A. ocellaris
A. percula
P. biaculeatus
Pomacentrus moluccensis
1/74.6/53.5
1/83.4/53.6
1/45.4/57.6
1/92.5/78.3
1/74.1/74.2
1/94.8/57.8
1/84.8/45.1
1/99.8/95
1/48/35
1/100/97
1/99.1/75.9
1/84.8/80.6
1/100/65
0.83/38.8/54.4
1/96.8/95.8
1/97.1/72.7
1/42/40
1/69.6/77.1
1/98.4/82.2
1/62.1/71.7
0.98/33.6/44
1/47.1/41.3
1/99.1/63
1/100/92.6
1/71.6/49.2
1/100/91
3.0

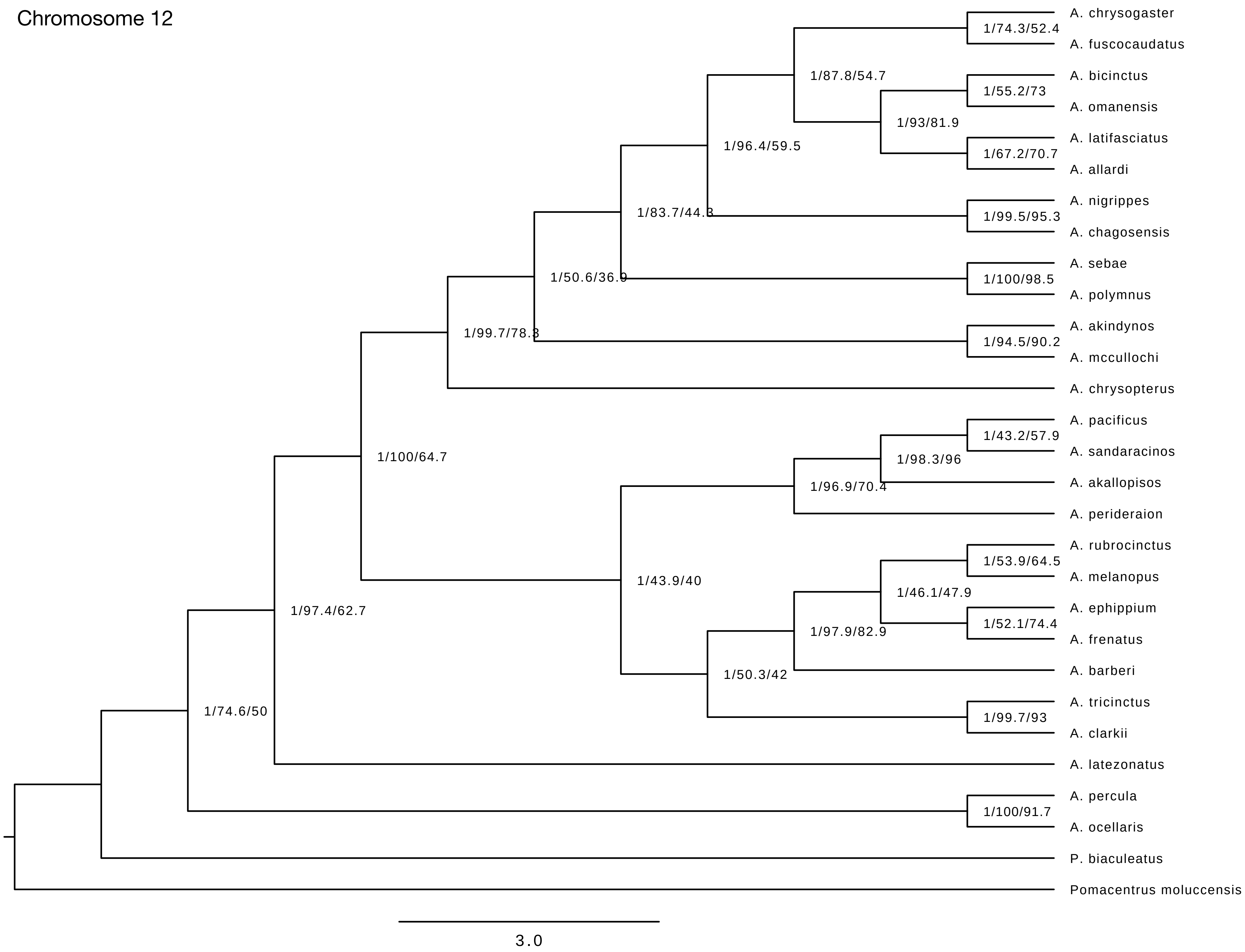
Chromosome 12
A. chrysogaster
A. fuscocaudatus
A. bicinctus
A. omanensis
A. latifasciatus
A. allardi
A. nigrippes
A. chagosensis
A. sebae
A. polymnus
A. akindynos
A. mccullochi
A. chrysopterus
A. pacificus
A. sandaracinos
A. akallopisos
A. perideraion
A. rubrocinctus
A. melanopus
A. ephippium
A. frenatus
A. barberi
A. tricinctus
A. clarkii
A. latezonatus
A. percula
A. ocellaris
P. biaculeatus
Pomacentrus moluccensis
1/74.3/52.4
1/87.8/54.7
1/55.2/73
1/93/81.9
1/67.2/70.7
1/96.4/59.5
1/83.7/44.3
1/99.5/95.3
1/50.6/36.9
1/100/98.5
1/99.7/78.3
1/94.5/90.2
1/100/64.7
1/43.2/57.9
1/98.3/96
1/96.9/70.4
1/53.9/64.5
1/43.9/40
1/46.1/47.9
1/97.4/62.7
1/52.1/74.4
1/97.9/82.9
1/50.3/42
1/99.7/93
1/74.6/50
1/100/91.7
3.0

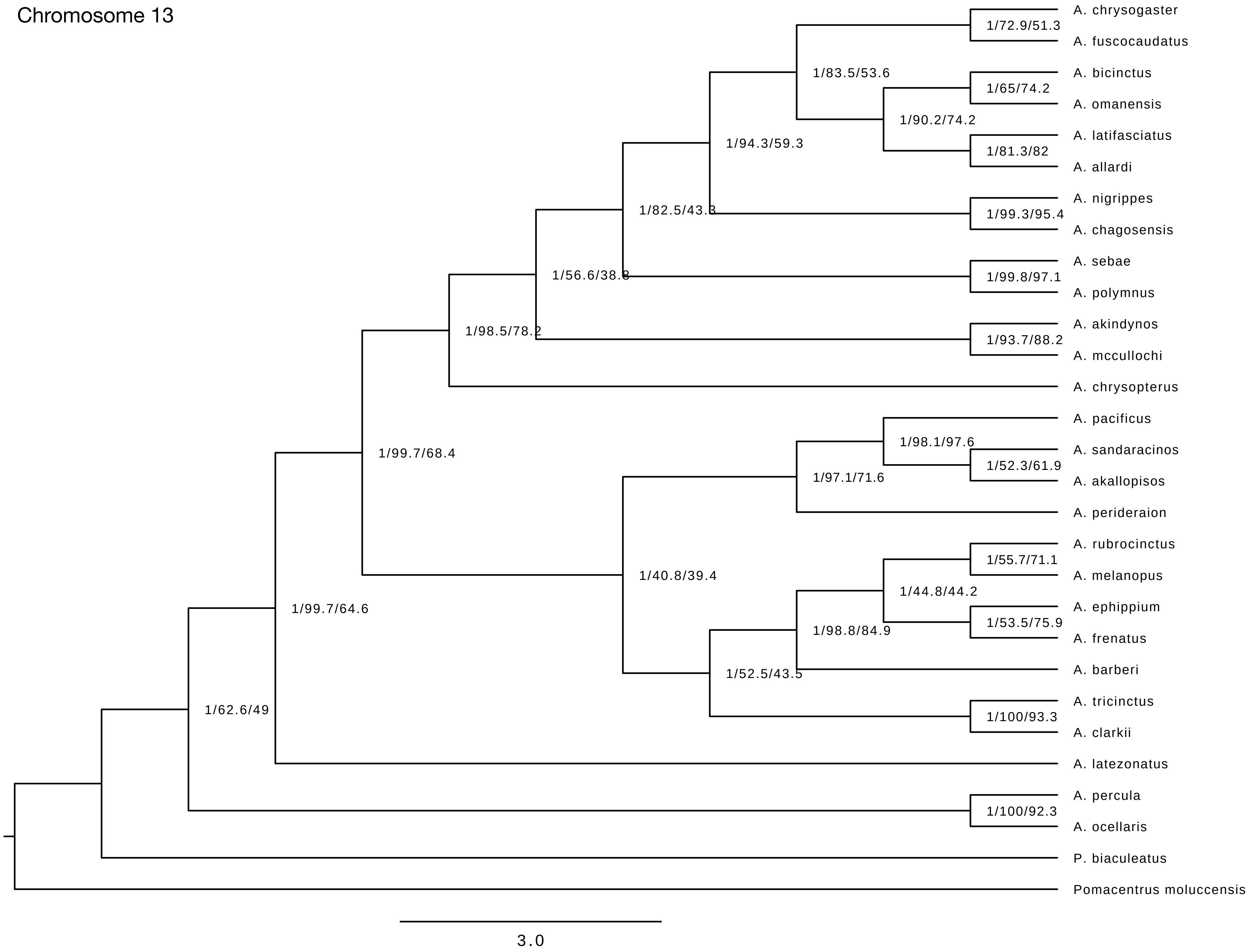
Chromosome 13
A. chrysogaster
A. fuscocaudatus
A. bicinctus
A. omanensis
A. latifasciatus
A. allardi
A. nigrippes
A. chagosensis
A. sebae
A. polymnus
A. akindynos
A. mccullochi
A. chrysopterus
A. pacificus
A. sandaracinos
A. akallopisos
A. perideraion
A. rubrocinctus
A. melanopus
A. ephippium
A. frenatus
A. barberi
A. tricinctus
A. clarkii
A. latezonatus
A. percula
A. ocellaris
P. biaculeatus
Pomacentrus moluccensis
1/72.9/51.3
1/83.5/53.6
1/65/74.2
1/90.2/74.2
1/81.3/82
1/94.3/59.3
1/82.5/43.3
1/99.3/95.4
1/56.6/38.8
1/99.8/97.1
1/98.5/78.2
1/93.7/88.2
1/99.7/68.4
1/98.1/97.6
1/52.3/61.9
1/97.1/71.6
1/40.8/39.4
1/55.7/71.1
1/44.8/44.2
1/53.5/75.9
1/98.8/84.9
1/99.7/64.6
1/52.5/43.5
1/100/93.3
1/62.6/49
1/100/92.3
3.0

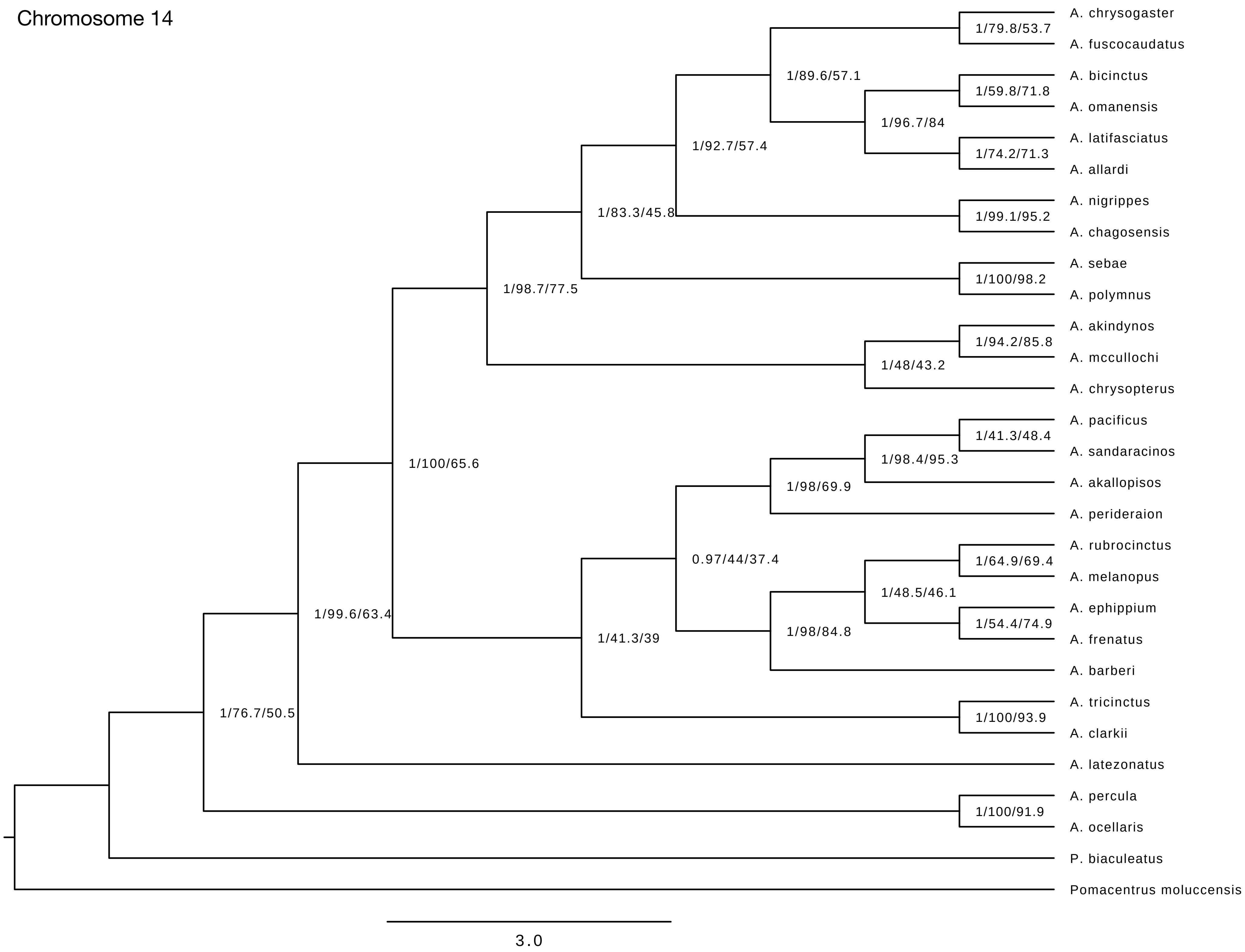
Chromosome 14
A. chrysogaster
A. fuscocaudatus
A. bicinctus
A. omanensis
A. latifasciatus
A. allardi
A. nigrippes
A. chagosensis
A. sebae
A. polymnus
A. akindynos
A. mccullochi
A. chrysopterus
A. pacificus
A. sandaracinos
A. akallopisos
A. perideraion
A. rubrocinctus
A. melanopus
A. ephippium
A. frenatus
A. barberi
A. tricinctus
A. clarkii
A. latezonatus
A. percula
A. ocellaris
P. biaculeatus
Pomacentrus moluccensis
1/79.8/53.7
1/89.6/57.1
1/59.8/71.8
1/96.7/84
1/74.2/71.3
1/92.7/57.4
1/83.3/45.8
1/99.1/95.2
1/98.7/77.5
1/100/98.2
1/94.2/85.8
1/48/43.2
1/100/65.6
1/41.3/48.4
1/98.4/95.3
1/98/69.9
0.97/44/37.4
1/64.9/69.4
1/48.5/46.1
1/54.4/74.9
1/99.6/63.4
1/98/84.8
1/41.3/39
1/76.7/50.5
1/100/93.9
1/100/91.9
3.0

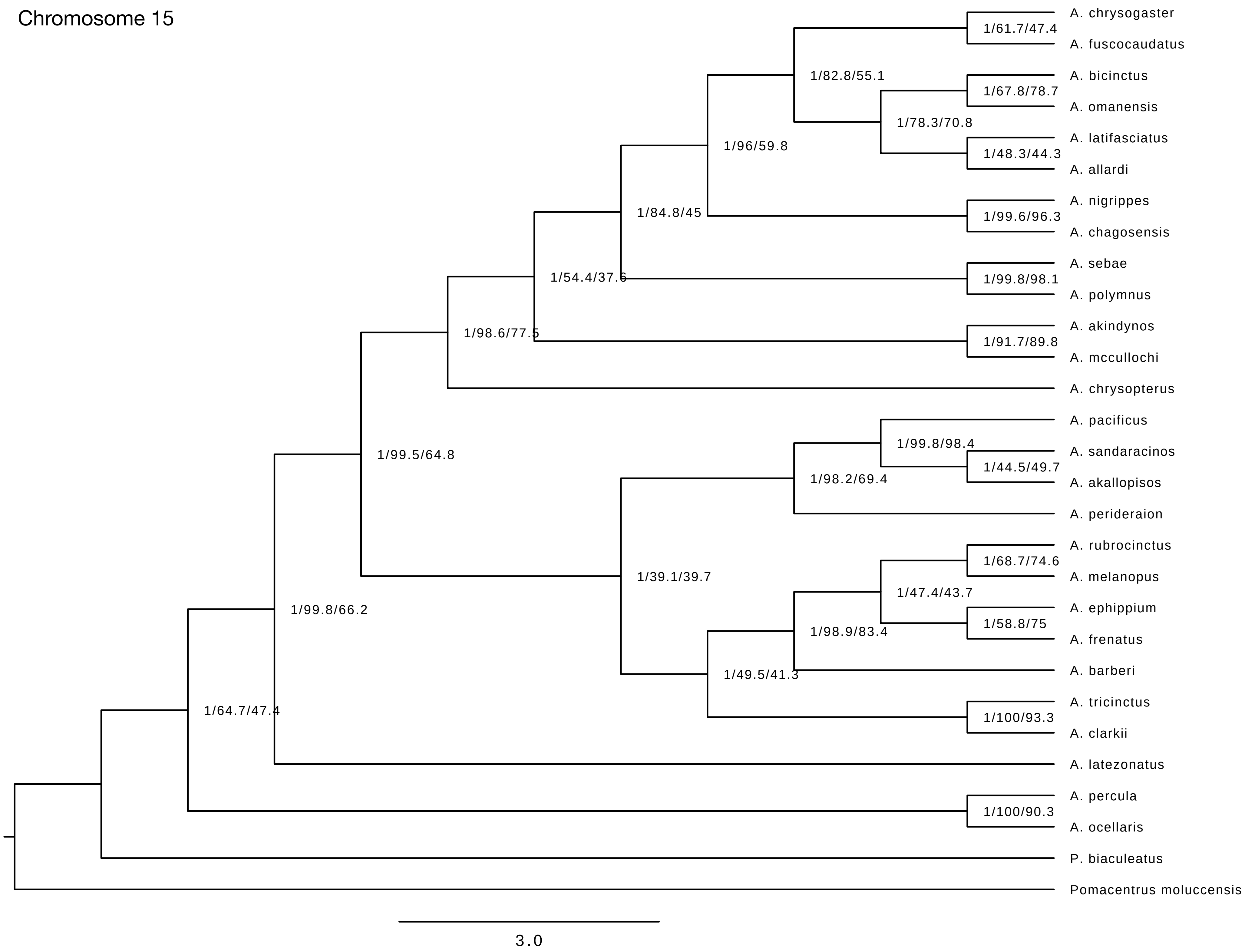
Chromosome 15
A. chrysogaster
A. fuscocaudatus
A. bicinctus
A. omanensis
A. latifasciatus
A. allardi
A. nigrippes
A. chagosensis
A. sebae
A. polymnus
A. akindynos
A. mccullochi
A. chrysopterus
A. pacificus
A. sandaracinos
A. akallopisos
A. perideraion
A. rubrocinctus
A. melanopus
A. ephippium
A. frenatus
A. barberi
A. tricinctus
A. clarkii
A. latezonatus
A. percula
A. ocellaris
P. biaculeatus
Pomacentrus moluccensis
1/61.7/47.4
1/82.8/55.1
1/67.8/78.7
1/78.3/70.8
1/48.3/44.3
1/96/59.8
1/84.8/45
1/99.6/96.3
1/54.4/37.6
1/99.8/98.1
1/98.6/77.5
1/91.7/89.8
1/99.5/64.8
1/99.8/98.4
1/44.5/49.7
1/98.2/69.4
1/39.1/39.7
1/68.7/74.6
1/47.4/43.7
1/58.8/75
1/98.9/83.4
1/49.5/41.3
1/100/93.3
1/99.8/66.2
1/64.7/47.4
1/100/90.3
3.0

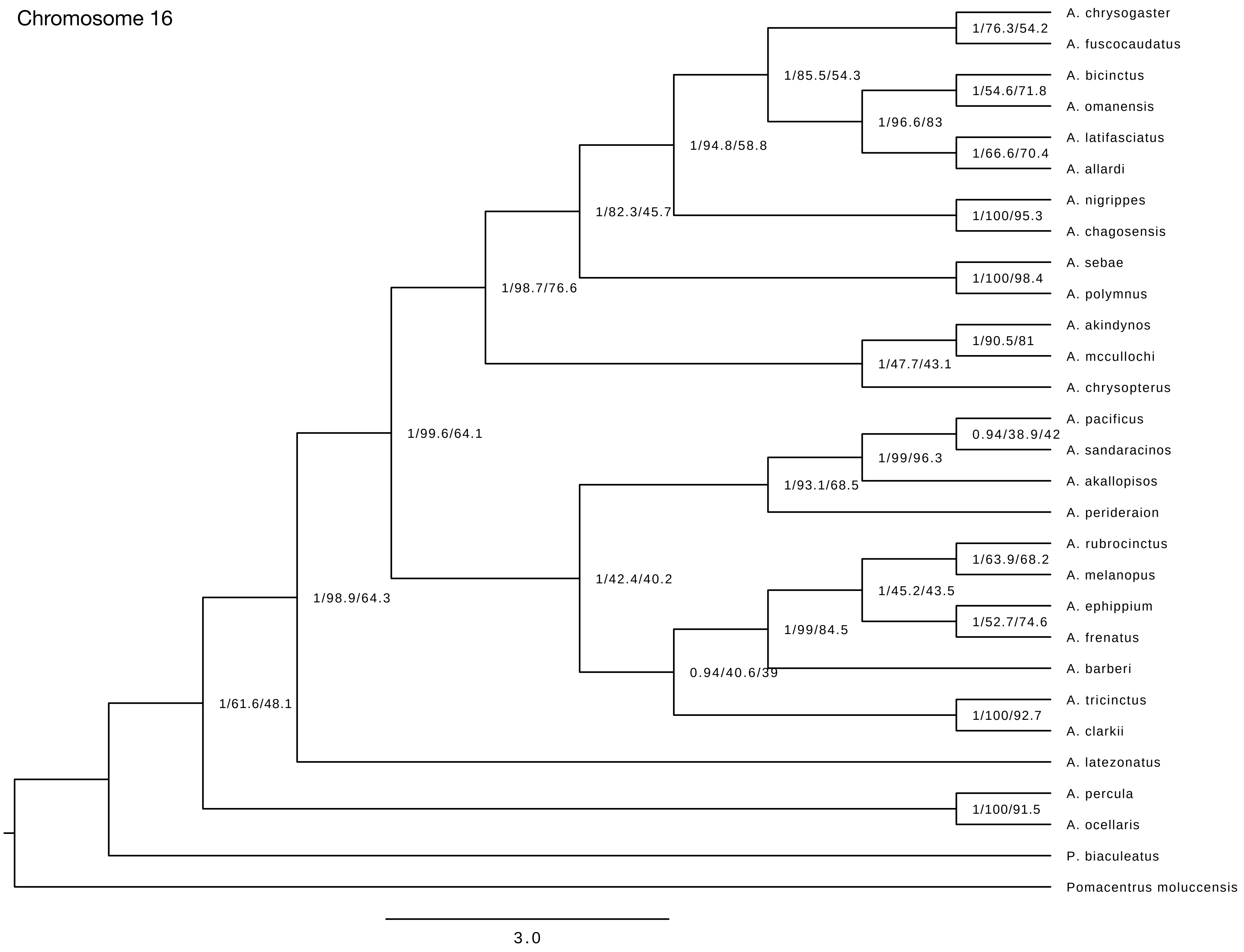
Chromosome 16
A. chrysogaster
A. fuscocaudatus
A. bicinctus
A. omanensis
A. latifasciatus
A. allardi
A. nigrippes
A. chagosensis
A. sebae
A. polymnus
A. akindynos
A. mccullochi
A. chrysopterus
A. pacificus
A. sandaracinos
A. akallopisos
A. perideraion
A. rubrocinctus
A. melanopus
A. ephippium
A. frenatus
A. barberi
A. tricinctus
A. clarkii
A. latezonatus
A. percula
A. ocellaris
P. biaculeatus
Pomacentrus moluccensis
1/76.3/54.2
1/85.5/54.3
1/54.6/71.8
1/96.6/83
1/94.8/58.8
1/66.6/70.4
1/82.3/45.7
1/100/95.3
1/98.7/76.6
1/100/98.4
1/90.5/81
1/47.7/43.1
1/99.6/64.1
0.94/38.9/42
1/99/96.3
1/93.1/68.5
1/63.9/68.2
1/42.4/40.2
1/45.2/43.5
1/98.9/64.3
1/52.7/74.6
1/99/84.5
0.94/40.6/39
1/61.6/48.1
1/100/92.7
1/100/91.5
3.0

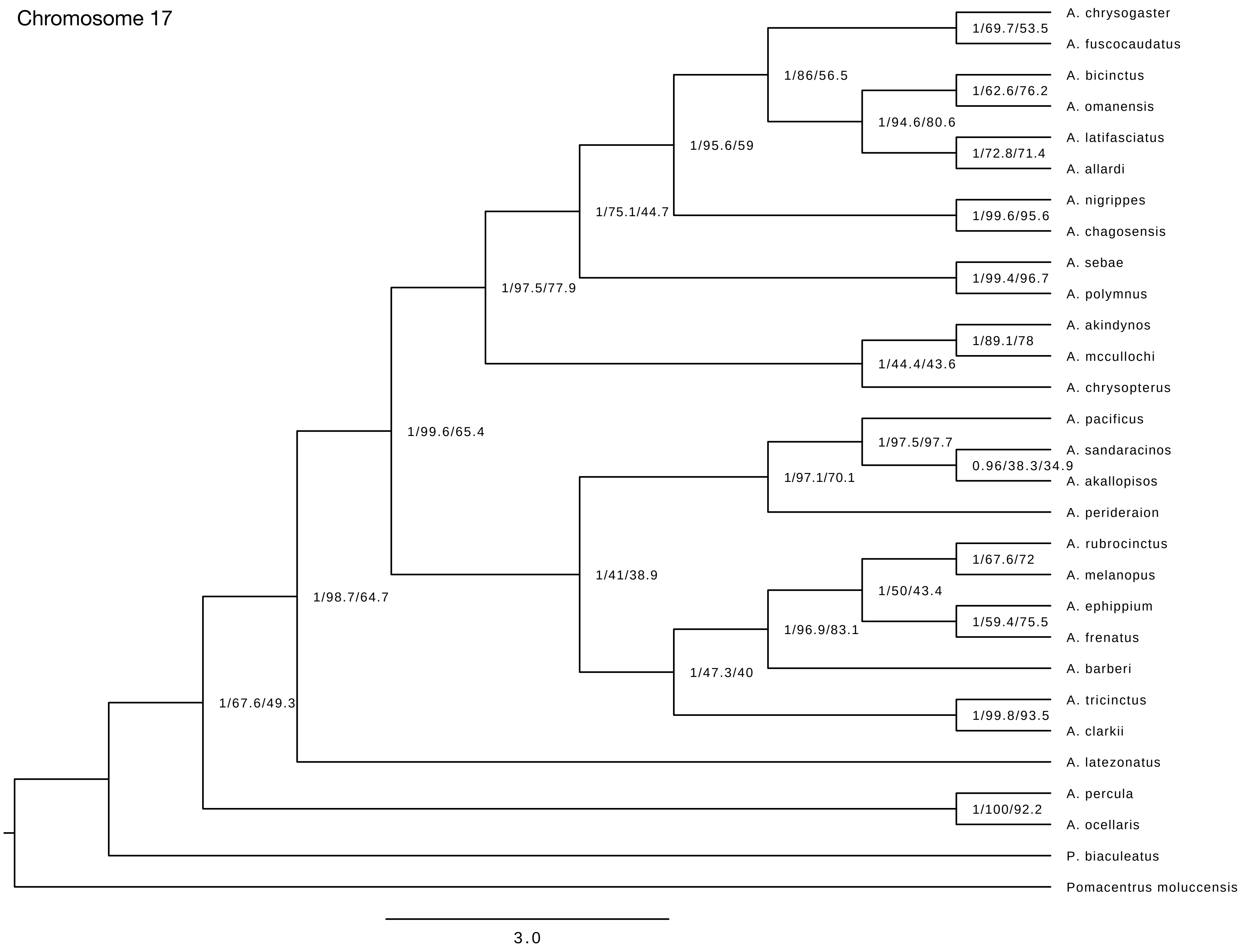
Chromosome 17
A. chrysogaster
A. fuscocaudatus
A. bicinctus
A. omanensis
A. latifasciatus
A. allardi
A. nigrippes
A. chagosensis
A. sebae
A. polymnus
A. akindynos
A. mccullochi
A. chrysopterus
A. pacificus
A. sandaracinos
A. akallopisos
A. perideraion
A. rubrocinctus
A. melanopus
A. ephippium
A. frenatus
A. barberi
A. tricinctus
A. clarkii
A. latezonatus
A. percula
A. ocellaris
P. biaculeatus
Pomacentrus moluccensis
1/69.7/53.5
1/86/56.5
1/62.6/76.2
1/94.6/80.6
1/72.8/71.4
1/95.6/59
1/75.1/44.7
1/99.6/95.6
1/97.5/77.9
1/99.4/96.7
1/89.1/78
1/44.4/43.6
1/99.6/65.4
1/97.5/97.7
0.96/38.3/34.9
1/97.1/70.1
1/67.6/72
1/41/38.9
1/50/43.4
1/98.7/64.7
1/59.4/75.5
1/96.9/83.1
1/47.3/40
1/67.6/49.3
1/99.8/93.5
1/100/92.2
3.0

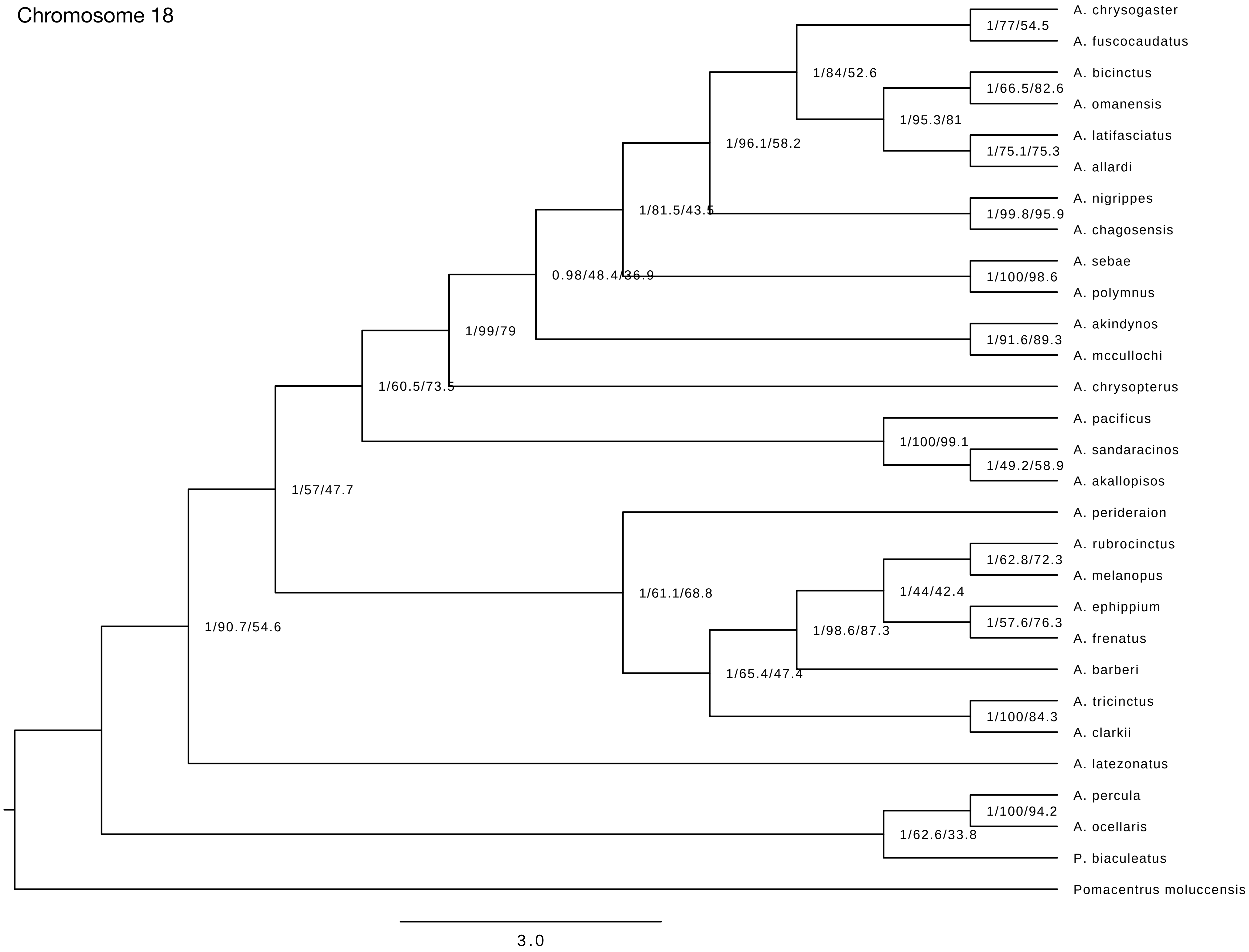

Chromosome 18
A. chrysogaster
A. fuscocaudatus
A. bicinctus
A. omanensis
A. latifasciatus
A. allardi
A. nigrippes
A. chagosensis
A. sebae
A. polymnus
A. akindynos
A. mccullochi
A. chrysopterus
A. pacificus
A. sandaracinos
A. akallopisos
A. perideraion
A. rubrocinctus
A. melanopus
A. ephippium
A. frenatus
A. barberi
A. tricinctus
A. clarkii
A. latezonatus
A. percula
A. ocellaris
P. biaculeatus
Pomacentrus moluccensis
1/77/54.5
1/84/52.6
1/66.5/82.6
1/95.3/81
1/75.1/75.3
1/96.1/58.2
1/81.5/43.5
1/99.8/95.9
0.98/48.4/36.9
1/100/98.6
1/99/79
1/91.6/89.3
1/60.5/73.5
1/100/99.1
1/49.2/58.9
1/57/47.7
1/62.8/72.3
1/61.1/68.8
1/44/42.4
1/57.6/76.3
1/90.7/54.6
1/98.6/87.3
1/65.4/47.4
1/100/84.3
1/100/94.2
1/62.6/33.8
3.0

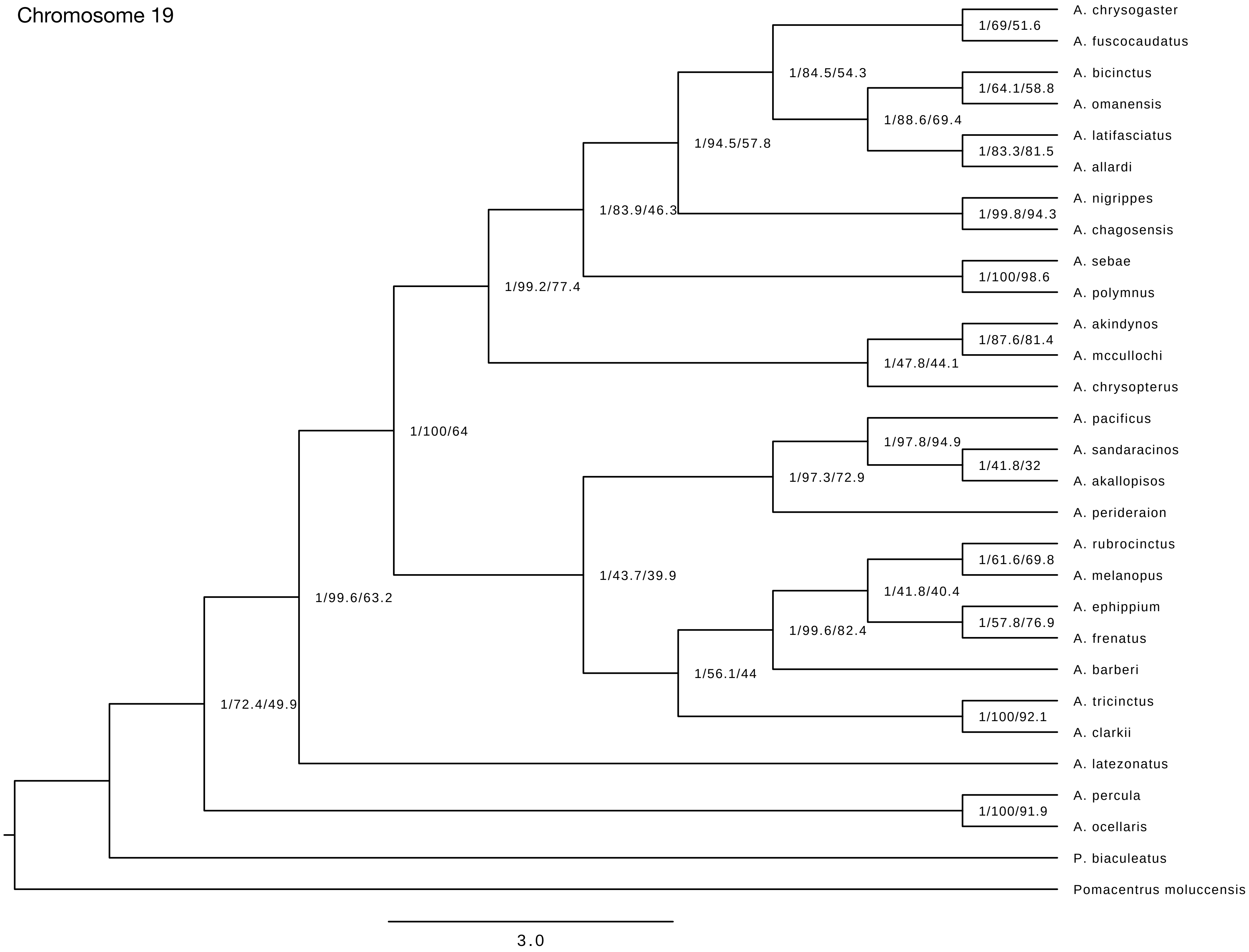
Chromosome 19
A. chrysogaster
A. fuscocaudatus
A. bicinctus
A. omanensis
A. latifasciatus
A. allardi
A. nigrippes
A. chagosensis
A. sebae
A. polymnus
A. akindynos
A. mccullochi
A. chrysopterus
A. pacificus
A. sandaracinos
A. akallopisos
A. perideraion
A. rubrocinctus
A. melanopus
A. ephippium
A. frenatus
A. barberi
A. tricinctus
A. clarkii
A. latezonatus
A. percula
A. ocellaris
P. biaculeatus
Pomacentrus moluccensis
1/69/51.6
1/84.5/54.3
1/64.1/58.8
1/88.6/69.4
1/83.3/81.5
1/94.5/57.8
1/83.9/46.3
1/99.8/94.3
1/100/98.6
1/99.2/77.4
1/87.6/81.4
1/47.8/44.1
1/100/64
1/97.8/94.9
1/41.8/32
1/97.3/72.9
1/61.6/69.8
1/43.7/39.9
1/41.8/40.4
1/99.6/63.2
1/57.8/76.9
1/99.6/82.4
1/56.1/44
1/72.4/49.9
1/100/92.1
1/100/91.9
3.0

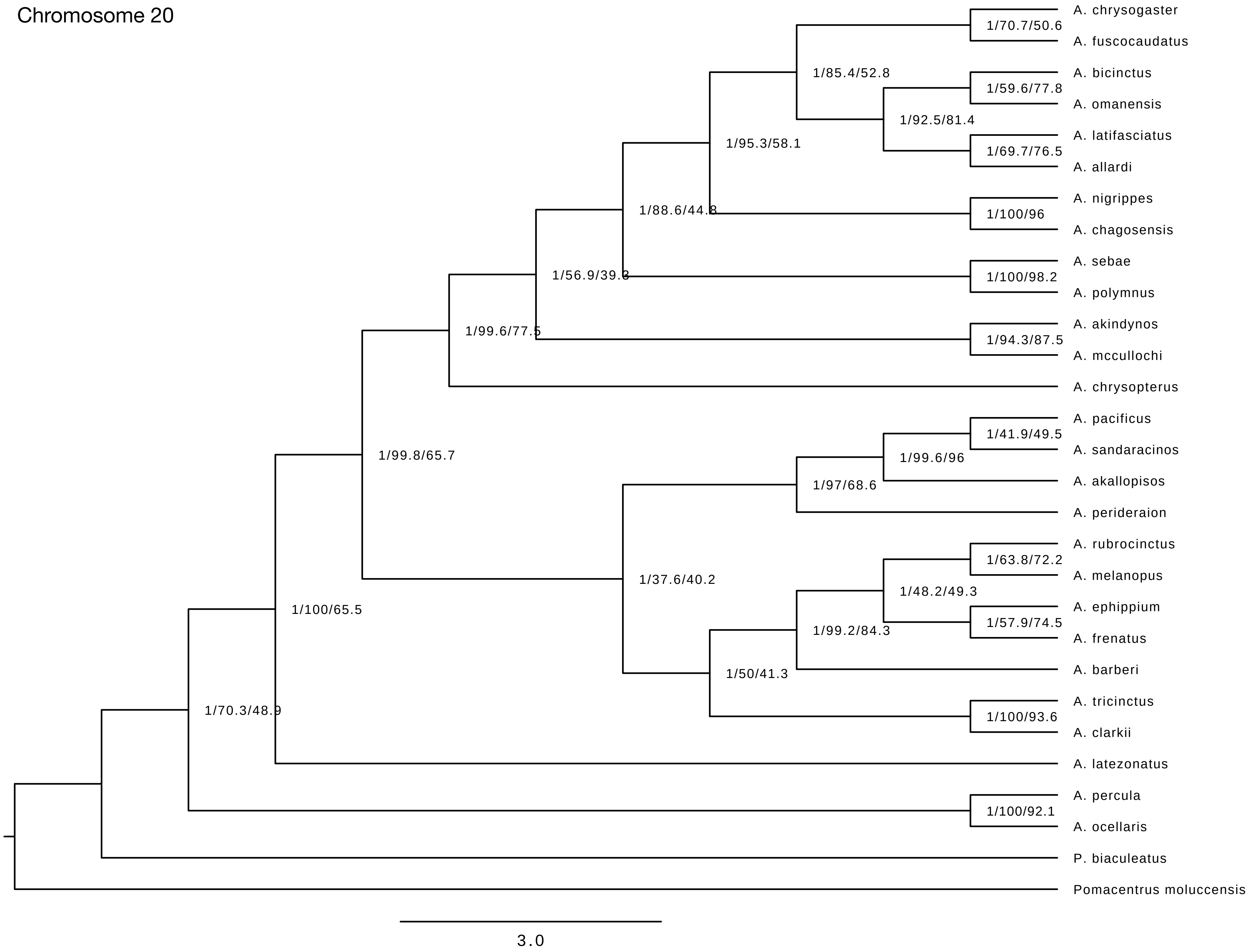
Chromosome 20
A. chrysogaster
A. fuscocaudatus
1/70.7/50.6
1/85.4/52.8
A. bicinctus
A. omanensis
1/59.6/77.8
1/92.5/81.4
A. latifasciatus
A. allardi
1/69.7/76.5
1/95.3/58.1
A. nigrippes
A. chagosensis
1/100/96
1/88.6/44.8
A. sebae
A. polymnus
1/100/98.2
1/56.9/39.3
A. akindynos
A. mccullochi
1/94.3/87.5
1/99.6/77.5
A. chrysopterus
1/99.8/65.7
A. pacificus
A. sandaracinos
1/41.9/49.5
1/99.6/96
A. akallopisos
1/97/68.6
A. perideraion
A. rubrocinctus
A. melanopus
1/63.8/72.2
1/48.2/49.3
A. ephippium
A. frenatus
1/57.9/74.5
1/99.2/84.3
A. barberi
1/37.6/40.2
1/50/41.3
A. tricinctus
A. clarkii
1/100/93.6
1/100/65.5
A. latezonatus
1/70.3/48.9
A. percula
A. ocellaris
1/100/92.1
P. biaculeatus
Pomacentrus moluccensis
3.0

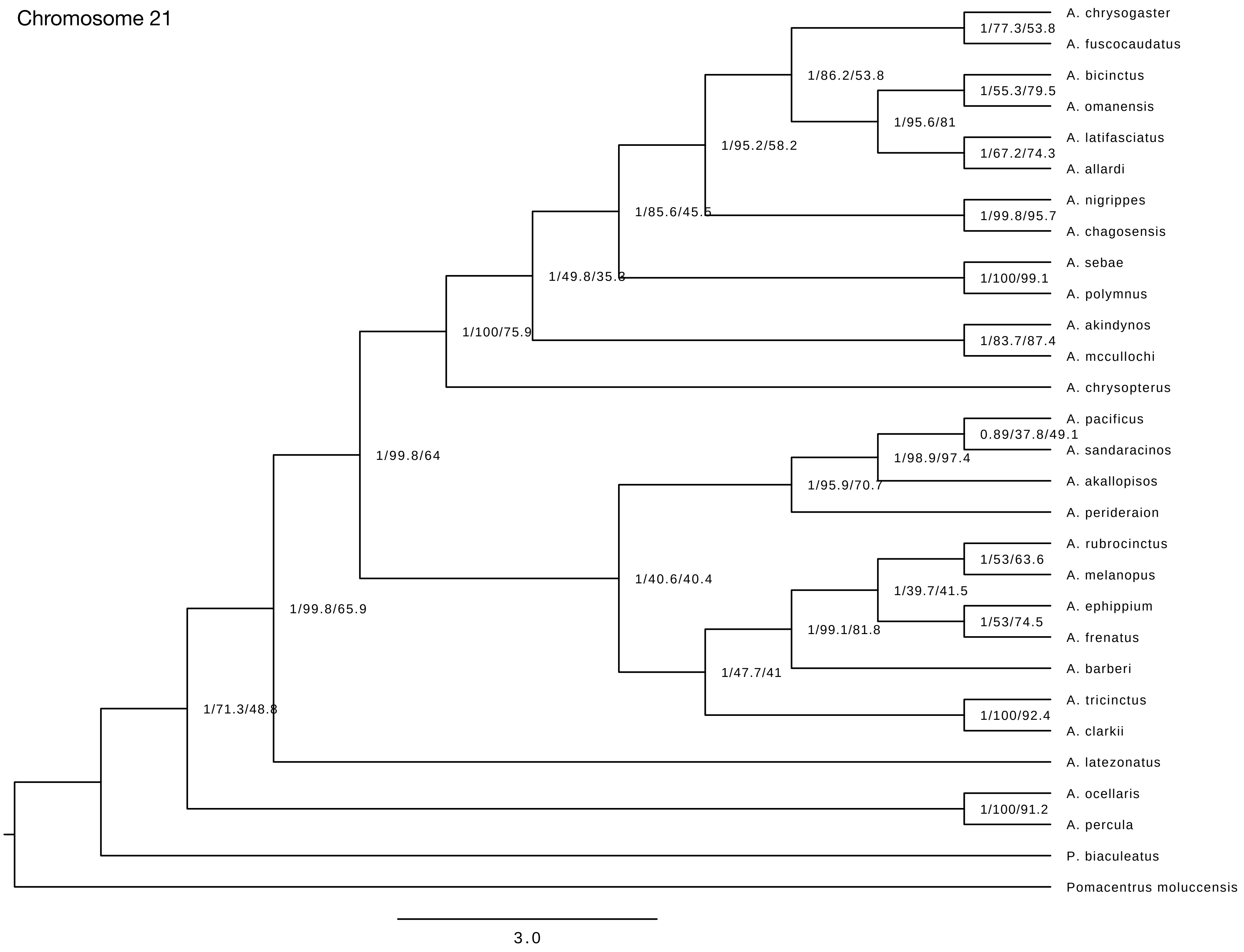
Chromosome 21
A. chrysogaster
A. fuscocaudatus
A. bicinctus
A. omanensis
A. latifasciatus
A. allardi
A. nigrippes
A. chagosensis
A. sebae
A. polymnus
A. akindynos
A. mccullochi
A. chrysopterus
A. pacificus
A. sandaracinos
A. akallopisos
A. perideraion
A. rubrocinctus
A. melanopus
A. ephippium
A. frenatus
A. barberi
A. tricinctus
A. clarkii
A. latezonatus
A. ocellaris
A. percula
P. biaculeatus
Pomacentrus moluccensis
1/77.3/53.8
1/86.2/53.8
1/55.3/79.5
1/95.6/81
1/67.2/74.3
1/95.2/58.2
1/99.8/95.7
1/85.6/45.5
1/100/99.1
1/49.8/35.3
1/100/75.9
1/83.7/87.4
1/99.8/64
0.89/37.8/49.1
1/98.9/97.4
1/95.9/70.7
1/53/63.6
1/39.7/41.5
1/40.6/40.4
1/53/74.5
1/99.8/65.9
1/99.1/81.8
1/47.7/41
1/100/92.4
1/71.3/48.8
1/100/91.2
3.0

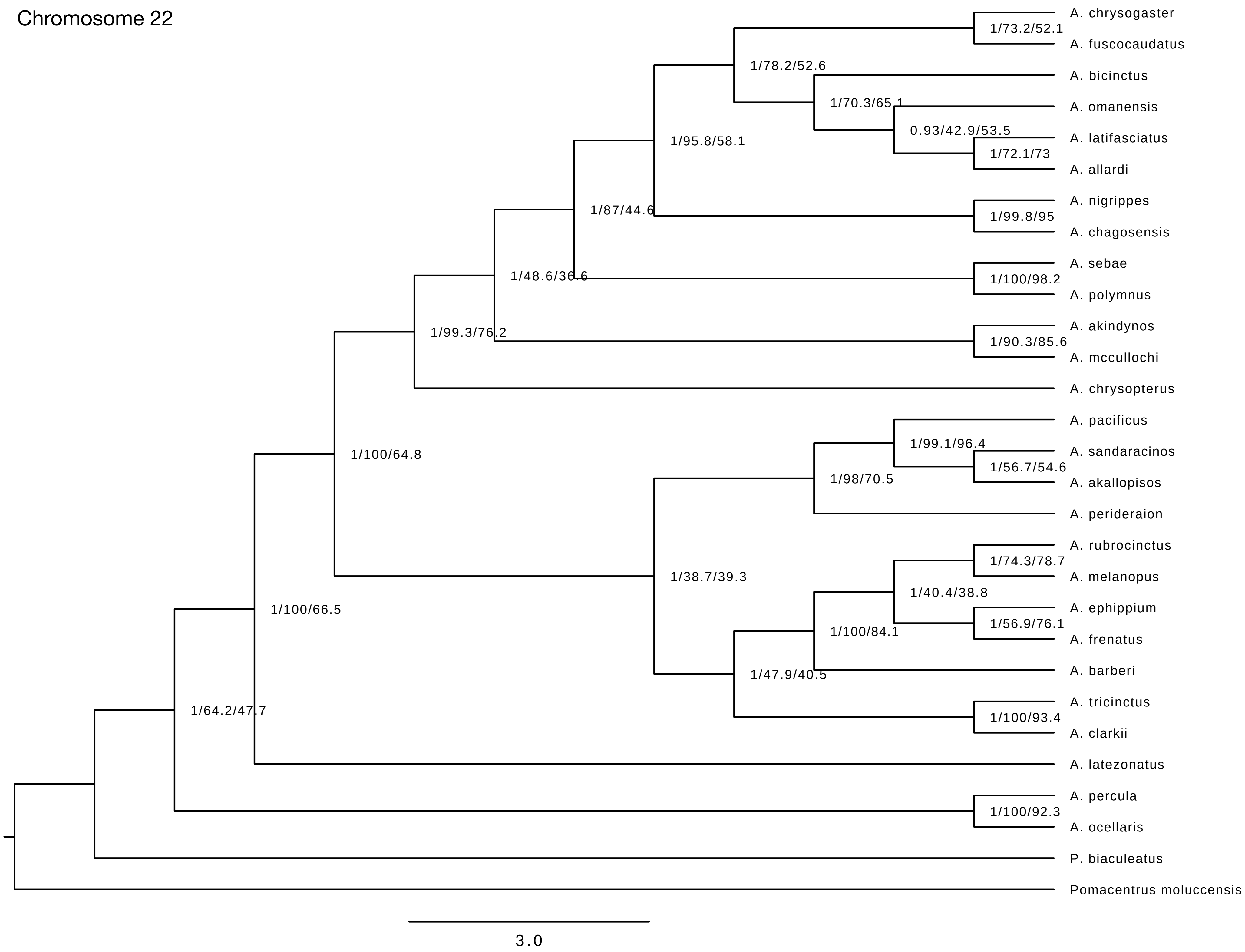

Chromosome 22
A. chrysogaster
A. fuscocaudatus
A. bicinctus
A. omanensis
A. latifasciatus
A. allardi
A. nigrippes
A. chagosensis
A. sebae
A. polymnus
A. akindynos
A. mccullochi
A. chrysopterus
A. pacificus
A. sandaracinos
A. akallopisos
A. perideraion
A. rubrocinctus
A. melanopus
A. ephippium
A. frenatus
A. barberi
A. tricinctus
A. clarkii
A. latezonatus
A. percula
A. ocellaris
P. biaculeatus
Pomacentrus moluccensis
1/73.2/52.1
1/78.2/52.6
1/70.3/65.1
0.93/42.9/53.5
1/72.1/73
1/95.8/58.1
1/87/44.6
1/99.8/95
1/48.6/36.6
1/100/98.2
1/99.3/76.2
1/90.3/85.6
1/100/64.8
1/99.1/96.4
1/56.7/54.6
1/98/70.5
1/38.7/39.3
1/74.3/78.7
1/40.4/38.8
1/56.9/76.1
1/100/84.1
1/47.9/40.5
1/100/93.4
1/100/66.5
1/64.2/47.7
1/100/92.3
3.0

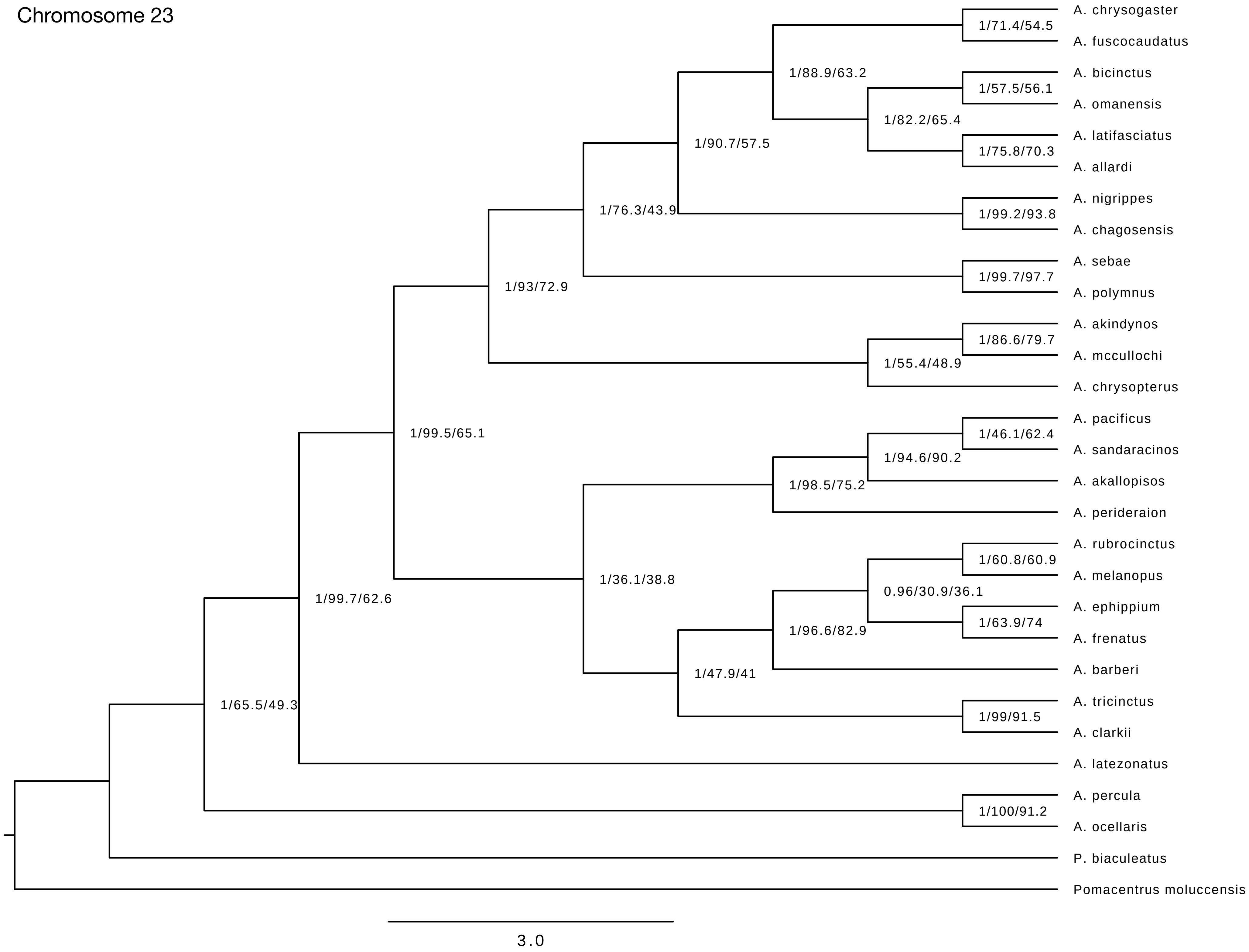
Chromosome 23
A. chrysogaster
A. fuscocaudatus
1/71.4/54.5
1/88.9/63.2
A. bicinctus
A. omanensis
1/57.5/56.1
1/82.2/65.4
A. latifasciatus
A. allardi
1/75.8/70.3
1/90.7/57.5
A. nigrippes
A. chagosensis
1/99.2/93.8
1/76.3/43.9
A. sebae
A. polymnus
1/99.7/97.7
1/93/72.9
A. akindynos
A. mccullochi
1/86.6/79.7
1/55.4/48.9
A. chrysopterus
A. pacificus
A. sandaracinos
1/46.1/62.4
1/94.6/90.2
A. akallopisos
1/98.5/75.2
A. perideraion
1/99.5/65.1
A. rubrocinctus
A. melanopus
1/60.8/60.9
0.96/30.9/36.1
A. ephippium
A. frenatus
1/63.9/74
1/96.6/82.9
A. barberi
1/36.1/38.8
1/47.9/41
A. tricinctus
A. clarkii
1/99/91.5
1/99.7/62.6
A. latezonatus
1/65.5/49.3
A. percula
A. ocellaris
1/100/91.2
P. biaculeatus
Pomacentrus moluccensis
3.0

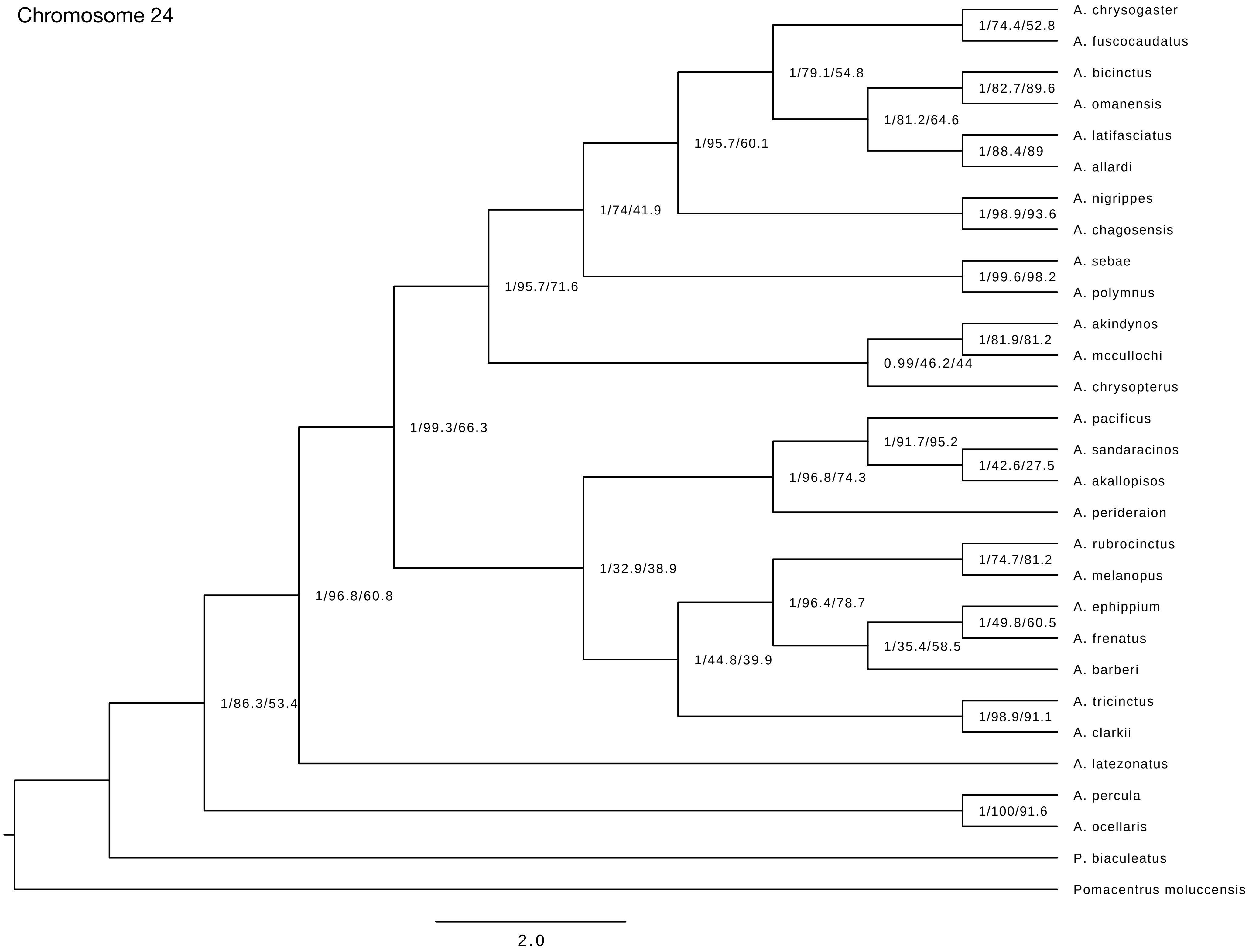
Chromosome 24
A. chrysogaster
A. fuscocaudatus
1/74.4/52.8
1/79.1/54.8
A. bicinctus
A. omanensis
1/82.7/89.6
1/81.2/64.6
A. latifasciatus
A. allardi
1/88.4/89
1/95.7/60.1
A. nigrippes
A. chagosensis
1/98.9/93.6
1/74/41.9
A. sebae
A. polymnus
1/99.6/98.2
1/95.7/71.6
A. akindynos
A. mccullochi
1/81.9/81.2
0.99/46.2/44
A. chrysopterus
1/99.3/66.3
A. pacificus
A. sandaracinos
A. akallopisos
1/91.7/95.2
1/42.6/27.5
1/96.8/74.3
A. perideraion
A. rubrocinctus
A. melanopus
1/74.7/81.2
1/32.9/38.9
1/96.4/78.7
A. ephippium
A. frenatus
1/49.8/60.5
1/35.4/58.5
A. barberi
1/44.8/39.9
A. tricinctus
A. clarkii
1/98.9/91.1
1/96.8/60.8
1/86.3/53.4
A. latezonatus
A. percula
A. ocellaris
1/100/91.6
P. biaculeatus
Pomacentrus moluccensis
2.0
